# Early Alzheimer’s disease pathology in human cortex is associated with a transient phase of distinct cell states

**DOI:** 10.1101/2023.06.03.543569

**Authors:** Vahid Gazestani, Tushar Kamath, Naeem M. Nadaf, SJ Burris, Brendan Rooney, Antti Junkkari, Charles Vanderburg, Tuomas Rauramaa, Martine Therrien, Matthew Tegtmeyer, Sanna-Kaisa Herukka, Abdulraouf Abdulraouf, Samuel Marsh, Tarja Malm, Mikko Hiltunen, Ralda Nehme, Beth Stevens, Ville Leinonen, Evan Z. Macosko

**Author notes:** These authors contributed equally to this work (Vahid Gazestani, Tushar Kamath).

## Abstract

Cellular perturbations underlying Alzheimer’s disease are primarily studied in human postmortem samples and model organisms. Here we generated a single-nucleus atlas from a rare cohort of cortical biopsies from living individuals with varying degrees of Alzheimer’s disease pathology. We next performed a systematic cross-disease and cross-species integrative analysis to identify a set of cell states that are specific to early AD pathology. These changes–which we refer to as the Early Cortical Amyloid Response—were prominent in neurons, wherein we identified a transient state of hyperactivity preceding loss of excitatory neurons, which correlated with the selective loss of layer 1 inhibitory neurons. Microglia overexpressing neuroinflammatory-related processes also expanded as AD pathological burden increased. Lastly, both oligodendrocytes and pyramidal neurons upregulated genes associated with amyloid beta production and processing during this early hyperactive phase. Our integrative analysis provides an organizing framework for targeting circuit dysfunction, neuroinflammation, and amyloid production early in AD pathogenesis.

## Introduction

The first pathological sign of AD in the human cortex is the gradual accumulation of amyloid beta plaques, followed by the appearance of gliosis, misfolded tau, and neurodegeneration. Of critical importance is understanding the coordinated activities of neurons and glia during the early phases of the disease that initiate this pathogenic cascade^1, 2^. Several postmortem single-cell studies have begun identifying disease-associated cellular changes in AD, particularly at later histopathological disease stages^3–8^. Inference from postmortem samples can be complicated by peri-mortem transcriptional responses to agonal state, cessation of blood flow, hypoxia, and neuronal atrophy. Prior cytological^9–11^ and transcriptional^12^ analyses demonstrate a marked decline, particularly in neurons, of cell health within two to four hours postmortem. Consequently, several fundamental questions related to the early stages of AD remain unanswered, including which cell types are perturbed the most, what molecular mechanisms are dysregulated in neuronal types of different cortical layers, and how these early perturbations contribute to the production of misfolded proteins and progression of pathology in the human brain.

We reasoned that a deep analysis of samples from living individuals harboring various extents of amyloid deposits could provide an opportunity to comprehensively capture veridical states associated with early-stage AD pathology. We performed single-nucleus RNA-sequencing (snRNA-seq) on a rare set of surgical biopsy samples obtained from patients undergoing ventriculoperitoneal shunt placement for treatment of suspected normal pressure hydrocephalus (NPH). In a study of 335 individuals, 44% of these biopsies contained amyloid beta (Aꞵ) plaques^13^ and, most importantly, longitudinal follow up of multiple cohorts has indicated the presence of Aꞵ within NPH biopsies is strongly associated with a decline in cognitive performance and a future clinical diagnosis of AD^13–15^, demonstrating these biopsies capture early AD pathology. To ensure that our insights were not restricted to a single cohort and were specific to AD, we further developed an accurate integrative analysis framework to incorporate published postmortem and mouse model single-cell datasets to construct a compendium of 2.4 million uniformly annotated cell profiles across diseases and species. The resulting analyses revealed what we collectively term the Early Cortical Amyloid Response (ECAR): a suite of consistent tissue changes in specific cell types that co-occur with the initial onset of brain amyloidosis.

## Results

### A single-nucleus atlas of human brain biopsies to identify AD pathological perturbations

To capture cellular perturbations in cortex of living individuals in response to AD pathology, we collected biopsies–frozen within five minutes of surgical excision to ensure fidelity of *in vivo* transcriptional states–from the frontal cortices (Brodmann areas 8 and 9) of 52 patients with NPH (Figure 1A). Histopathological examination of the biopsies identified 19 with Aꞵ plaques (Aꞵ+), eight with both Aꞵ plaques and phosphorylated tau pathology (Aꞵ+Tau+), and 25 biopsies that had neither histopathology (Table S1). From the stereotactic position of the catheter insertion site recorded by post-surgical CT or MRI (Figure 1A), we determined that the anatomical location of sampling did not correlate with AD histopathological burden (Figure S1A). We further divided the Aꞵ+ biopsies into three groups by their level of plaque burden (Figure S1B). The extent of Aꞵ plaque and tau tangle burden within the biopsies correlated inversely with these patients’ Aꞵ-42 CSF levels (p-value < 0.001; Figure 1B), and positively with CSF levels of phosphorylated tau (p-value < 0.005; Figure 1B), consistent with prior biomarker studies of AD progression^16, 17^. Moreover, the CSF levels of phosphorylated tau in Aꞵ+ individuals were similar to individuals without histopathology (p-value >0.95; Student’s t-test) and significantly less than an independent cohort of 36 clinically diagnosed AD individuals (p-value < 0.004; Student’s t-test; Figure 1B). Collectively, these results suggest the severity of biopsy histopathology is representative of the overall burden in the brain.

**Figure 1.**
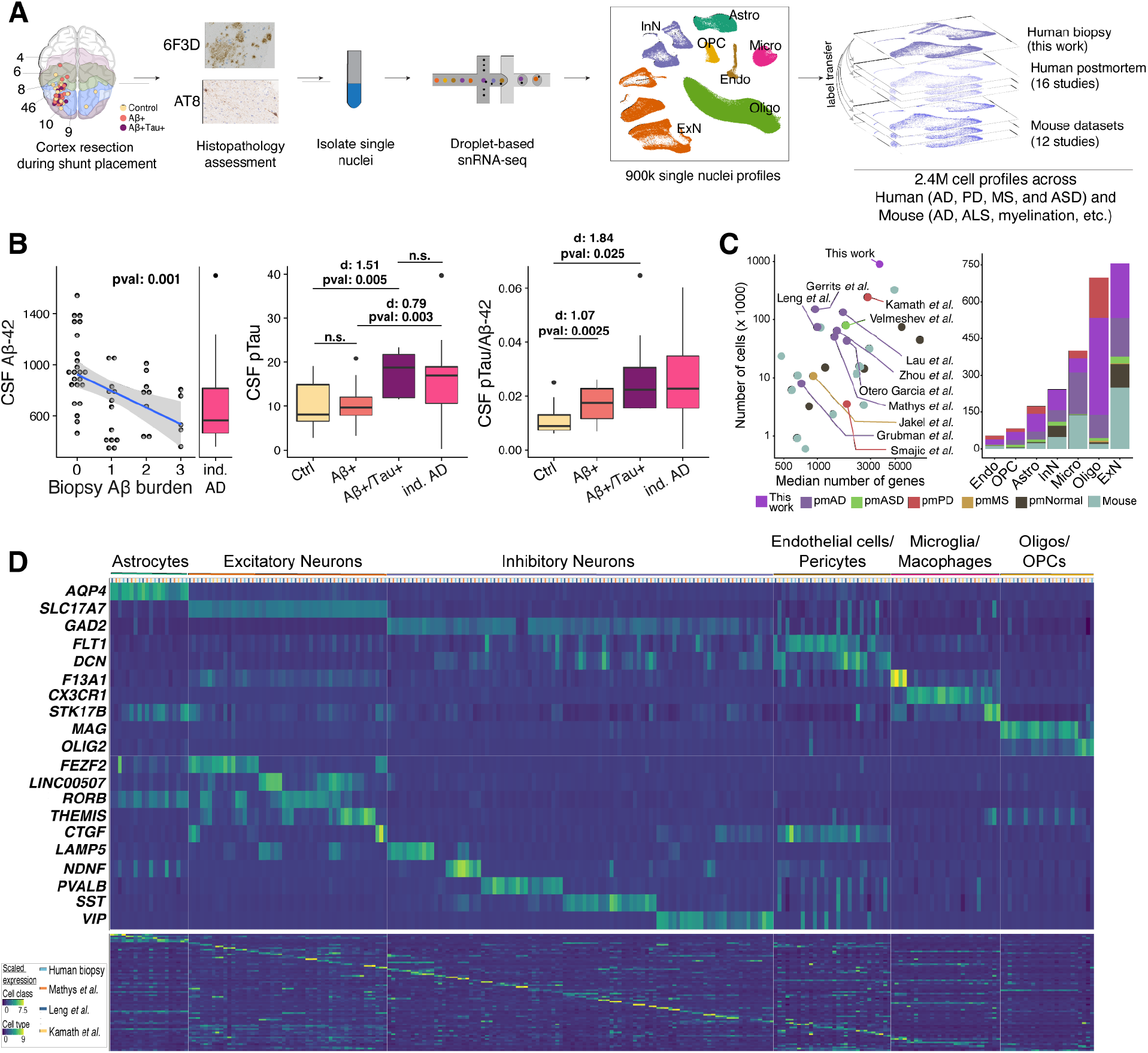
A fresh-tissue atlas of cortical states associated with AD pathology. ***A)** Schematic of the frontal cortex brain biopsy sampling workflow. Samples were stained and quantitatively assessed for AD histopathology by the 6F3D (Aꞵ) and AT8 (phosphorylated tau) antibodies (Methods). Brodmann areas are color-coded in the first panel. **B)** CSF Aꞵ-42 (left), phosphorylated tau (middle) and ratio of the two (right) in association with Aꞵ and tau burden scores (see Methods) in 49 subjects sampled. We have excluded three individuals for whom the CSF measurements were missing. The “ind. AD” refers to an independent cohort of 36 NPH patients who were clinically diagnosed with AD prior to, or within one year after, CSF collection. Cohen d (d) effect sizes are reported. **C)** A summary of datasets included in the integrative analysis. Case-control datasets of human brain diseases are labeled. pm, postmortem; ASD, autism spectrum disorder; PD, Parkinson’s disease; MS, multiple sclerosis. **D)** Expression of markers of cell classes (top), main neuronal classes (middle), and individual cell types (bottom) across four human studies of neurodegenerative disease from the integrative analysis. Each row indicates the normalized expression level of each gene across the select human postmortem datasets (color-coded on y-axis) and 82 cell types. A detailed analysis of cell types and associated markers can be found in Table S3.*

To explore the cell-type-specific changes associated with Aꞵ and tau histopathology in the cortex, we obtained 892,828 high-quality nuclei profiles from this biopsy cohort, with a median of 17,082 nuclei per individual. By unsupervised clustering^18^ (Methods), we identified the seven major classes of cells in the cortex: excitatory neurons (ExN; 222,449 nuclei), inhibitory neurons (InN; 83,702 nuclei), microglia (Micro; 59,624 nuclei), astrocytes (Astro; 73,487 nuclei), endothelial cells/pericytes (Endo; 22,407 nuclei), oligodendrocytes (Oligo; 396,292 nuclei), and oligodendrocyte progenitor cells (OPC; 34,867 nuclei) (Figure 1A). To increase our resolution, we repeated our clustering analysis within each class to identify a total of 82 cell types with a median size of 3,586 nuclei per type.

### An integrative analysis of biopsy and postmortem brain

Several studies have profiled brain cells under normal and disease conditions using human postmortem or mouse samples. However, a direct comparison of their results have been hampered by differences in sample qualities, dataset sizes, analysis pipelines, and cell type annotations. We reasoned that the size and quality of our biopsy dataset would be sufficiently analytically powered to conduct a comprehensive integrative analysis of these datasets with highly granular cell type specificity. For our integrative analysis, we considered 27 published single-cell/nuclei studies of the brain derived from both human disease studies and mouse disease models (Figure 1C). Human studies included postmortem samples from individuals with AD, Parkinson’s disease (PD), multiple sclerosis (MS), and autism spectrum disorder (ASD) (Table S2). Mouse datasets included models of AD and ALS, as well as de/re-myelination, aging, prenatal, and food deprivation conditions among others (Table S2). To accurately combine these datasets with our biopsy cohort, we developed an optimized single-cell integration framework that efficiently handled the substantial technical (e.g., sample preparation, sequencing platforms and depth) and biological (e.g., human vs mouse) variation that exists among these datasets (Methods). We employed three criteria to validate the quality of our integrative analysis results: 1) uniform mixing of the datasets across clusters; 2) cells expressing similar cortical cell type markers are aligned with each other across datasets and organisms; 3) reported cell type identities in each of the studies are preserved in the aligned space. A total of 2,406,980 cells were included in our integrative analysis after removing artifacts, low quality cells, and doublets. We next implemented a random walk method to transfer cell type annotations from our biopsy cohort to each of 27 other studies, thereby uniformly annotating all datasets to the 82 cell types (Figure 1D; Table S3). Comparison across human datasets demonstrated our biopsy cohort had among the highest number of cells sampled per cell type and minimal expression of artifactual genes often associated with sample quality and dissociation methods^19, 20^ (Figures S2-S4). Congruently, our attempts failed to achieve similarly high integration resolution after exclusion of our biopsy dataset from the analysis (data not shown).

We then investigated how agonal states and the postmortem interval affected gene expression patterns in different cell types by comparing our biopsy dataset with postmortem data, and identifying recurrent correlates with postmortem interval across datasets. Our analysis demonstrated a small but statistically significant decrease in gene expression levels in both excitatory (p-value < 0.038; Meta analysis) and inhibitory neurons (p-value < 0.024; Meta analysis, Figure S5A), as well as a trend towards increased gene expression levels in microglia (Figure S5A). Consistently, the ratio of glial to neuronal gene expression was lowest in the biopsy dataset, and this ratio increased with longer postmortem intervals within postmortem datasets (Figures S5B-S5D). Together, our results, in combination with the expression patterns of artifact associated genes (Figures S3 and S4), indicate loss of transcriptional complexity in both inhibitory and excitatory neurons, as well as an increase in artifact-related genes in microglial cells in response to peri- and post-mortem events.

### Meta-analysis reveals cortical cellular changes in early AD pathology

To identify cortical tissue changes across progression of AD pathology, we divided our biopsy samples into those with only Aꞵ pathology (Aꞵ+) and those with both Aꞵ and tau pathology (Aꞵ+Tau+). In parallel, we also analyzed two AD postmortem studies^3, 4^ that sampled both neuronal and glial cells from subjects with low Braak pathology staging and one dataset that only measured glia^6^ (Methods). We first tested for alterations in relative abundance of cell populations with increasing histopathological burden. A meta-analysis of cell proportions identified two neuron types–NDNF-PROX1 and LINC00507-COL5A2–that were significantly depleted (FDR-adjusted p-value < 0.05) (Figure 2A) in each of the cohorts with early amyloid pathology. The NDNF-PROX1 population expressed *NDNF* and *RELN*, markers of an interneuron type known to reside primarily in layer 1 (L1) of cortex^21^. The LINC0050s7-COL5A2 population expressed *CUX2* and *LINC00507,* consistent with a layer 2/3 (L2/3) telencephalic identity^21^. Seven additional cell types showed a trend toward significant loss (0.05 < FDR-adjusted p-value < 0.12; Table S4): two upper layer excitatory types (RORB-SCTR, LINC00507-ACVR1C), three inhibitory types (VIP-HTR3A and SST-PENK that are upper-layer-enriched,and VIP-NPSR1 that spans cortical layers; Figure S6A), one microglia type (CX3CR1) and one oligodendrocyte type (BACE2-L3MBTL4), while one microglia type (GPNMB-LPL) showed a trend toward expansion (Figure 2A). We further confirmed that the observed changes in neuronal populations do not correlate with the severity of any iNPH symptoms within the biopsy cohort (Figure S6B). Most importantly, our refined integration strategy and meta-analysis revealed similar alterations in neuronal and microglial proportions within each of the published postmortem AD case-control cohorts (Figure 2B), underscoring the robustness of the observed cellular changes associated with early-stage AD pathology.

**Figure 2.**
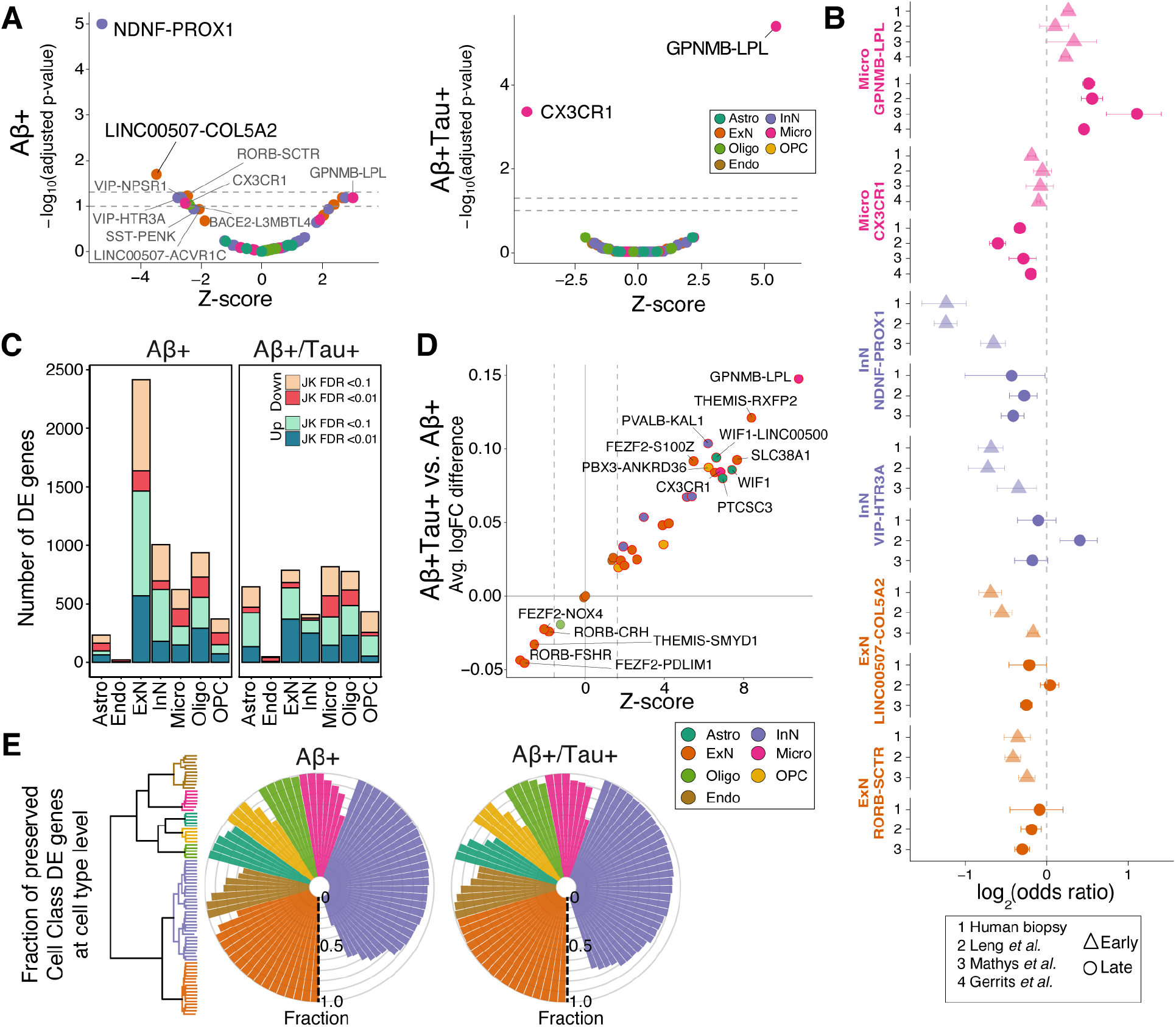
Identification of early- and late-stage cellular perturbations in AD. ***A)** Volcano plot of a meta-analysis of cell type proportional changes (Methods) in early- and late-stage AD-related samples. Cell types reaching significance are labeled. Colors indicate cell class assignment. Dashed lines represent FDR thresholds of 0.05 and 0.1. **B)** Individual log-odds ratios of six significant cell types in Aꞵ+ (triangles) and Aꞵ+Tau+ samples (circles) for our biopsy cohort and published postmortem AD case-control datasets. Whiskers indicate standard errors. **C)** Number of DE genes in each cell class, stratified by biopsy histopathology. JK: Jack-knife. **D)** Fold change pattern concordance of DE genes between Aꞵ+ and Aꞵ+Tau+ samples. The y-axis shows the average logFC difference between Aꞵ+Tau+ and Aꞵ+. The Z-scores on x-axis are based on the transformation of p-values from a paired t-test analysis on the union of top 300 protein-coding genes (sorted by their jack-knifed p-value) from each condition. **E)** Fraction of DE genes in Aꞵ+ and Aꞵ+Tau+ biopsies that are similarly up- or down-regulated between the seven major cell classes and their associated subtypes in biopsy samples. The fraction was calculated by examining the top 300 protein-coding DE genes at the cell class. The dendrogram illustrates the subdivision of the seven major cell classes to a total of 82 subtypes*.

In subjects with high histopathological burden, the proportional losses of the NDNF-PROX1 and LINC00507-COL5A2 neuronal populations were no longer significant (Figures 2A and 2B), likely due to additional loss of other cortical neurons, since the overall proportions of excitatory and inhibitory neurons were both lower in these subjects across cohorts (Figure S6C). Instead, we identified a significant (FDR-adjusted p-value < 0.05) expansion of the GPNMB-LPL microglia type and loss of the major homeostatic microglia population, marked by expression of *CX3CR1* (Figures 2A and 2B). In addition, while not consistently altered in all three human studies (meta-analysis p-value < 0.11), one astrocyte population expressing *CHI3L1* and *GFAP* increased in abundance in the late stages of disease in our biopsy cohort (p-value < 0.05; odds ratio (OR): 1.5), one postmortem cohort^4^ (p-value < 0.075; OR: 1.6), and in a mouse model of AD^22^ (p-value < 0.02; OR: 1.4) (Table S4). Together, these results indicate that gliosis becomes an increasingly prominent feature of cortical tissue as histopathology worsens.

Next, we examined how the transcriptional phenotype of each cell type changes across early and late histopathological stages of AD. We used a pseudocell-based strategy, coupled with mixed-effect modeling and jack-knifing (Methods), to robustly identify differentially expressed (DE) genes in both the Aꞵ+ and the Aꞵ+Tau+ biopsies (Figure 2C; Table S5). To better understand the association of gene perturbations with progression of AD pathology, we developed a metric to quantify the relative magnitude of transcriptional alteration across each cell type in early-versus late-stage samples (Methods). For most cell types and most notably in microglia populations, we found that the transcriptional alterations quantified in Aꞵ+Tau+ biopsies were consistent with, but stronger than, those changes measured in Aꞵ+ samples (Figure 2D). However, several excitatory neuron populations showed transcriptional perturbations in the Aꞵ+ samples that were absent in the Aꞵ+Tau+ biopsies (Figures 2C and 2D), indicating their passage through a distinct transcriptional state early in histopathological progression. To further assess the extent of overlap in dysregulated transcriptional programs among related cell populations, we calculated the fraction of DE genes in each of the seven major cell classes that show consistent DE within each of their constituent cell types. This comparison demonstrated that DE genes identified by the analysis of each of seven major cell classes exhibited highly preserved perturbation patterns (i.e., similar up- or down-regulation patterns) within their related cell types (Figures 2E and S6D). Collectively, our DE analyses demonstrated that: a) perturbation of the transcriptomes increases in magnitude as neuropathology worsens–with the exception of excitatory neurons, which show a distinct early phase response; and b) that individual cell type responses are largely similar within a major cell class.

### Neuronal loss and hyperactivity in early AD pathology

AD is increasingly recognized as a systems disease where interactions of different cell types define its pathological course. The strongest proportional change in our meta-analysis of the early AD pathological stage was the loss of NDNF-PROX1 inhibitory neurons (Figure 2A). We therefore wondered whether the loss of these inhibitory neurons could contribute to the onset and early progression of AD by induction of specific transcriptional states in other cortical cell types. To examine this, we correlated the fraction of NDNF-PROX1 inhibitory neurons with the extent of molecular perturbations in all other cell types (Methods). Intriguingly, applying this analysis to the Aꞵ+ biopsy samples identified a specific and significant (FDR-adjusted p-value < 0.01) correlation between NDNF-PROX1 depletion and upregulated ExN DE genes in the LINC00507-COL5A2 excitatory neurons (Figures 3A and S7A), which themselves are vulnerable to loss in early AD pathology (Figure 2A). Alternative analysis methods and robustness analyses confirmed the strength of association between the ExN DE signature in LINC00507-COL5A2 with the loss of *NDNF*+ expressing cells in Aꞵ+ individuals (Figures S7B-S7E). Moreover, this association was also significant (FDR-adjusted p-value < 0.05) within the control biopsy samples (Figures S7F and S7G), reinforcing that this pair of neuronal changes–loss of NDNF-PROX1 and induction of a specific transcriptional state in LINC00507-COL5A2–occurs early in disease. We next tested whether transcriptional response in LINC00507-COL5A2 neurons is specifically induced by loss of NDNF-PROX1 cells, or is also associated with the loss of other inhibitory neurons. Importantly, the relationship between the inhibitory neuron loss and ExN transcriptional state was specific to NDNF-PROX1 and VIP-HTR3A inhibitory neurons in Aꞵ+ biopsies (Figure 3B), the two most depleted inhibitory cell types in the early stage of AD. These results suggest a tight and specific coupling between layer 1 inhibitory neuron loss and transcriptional alterations in layer 2/3 excitatory neurons.

**Figure 3.**
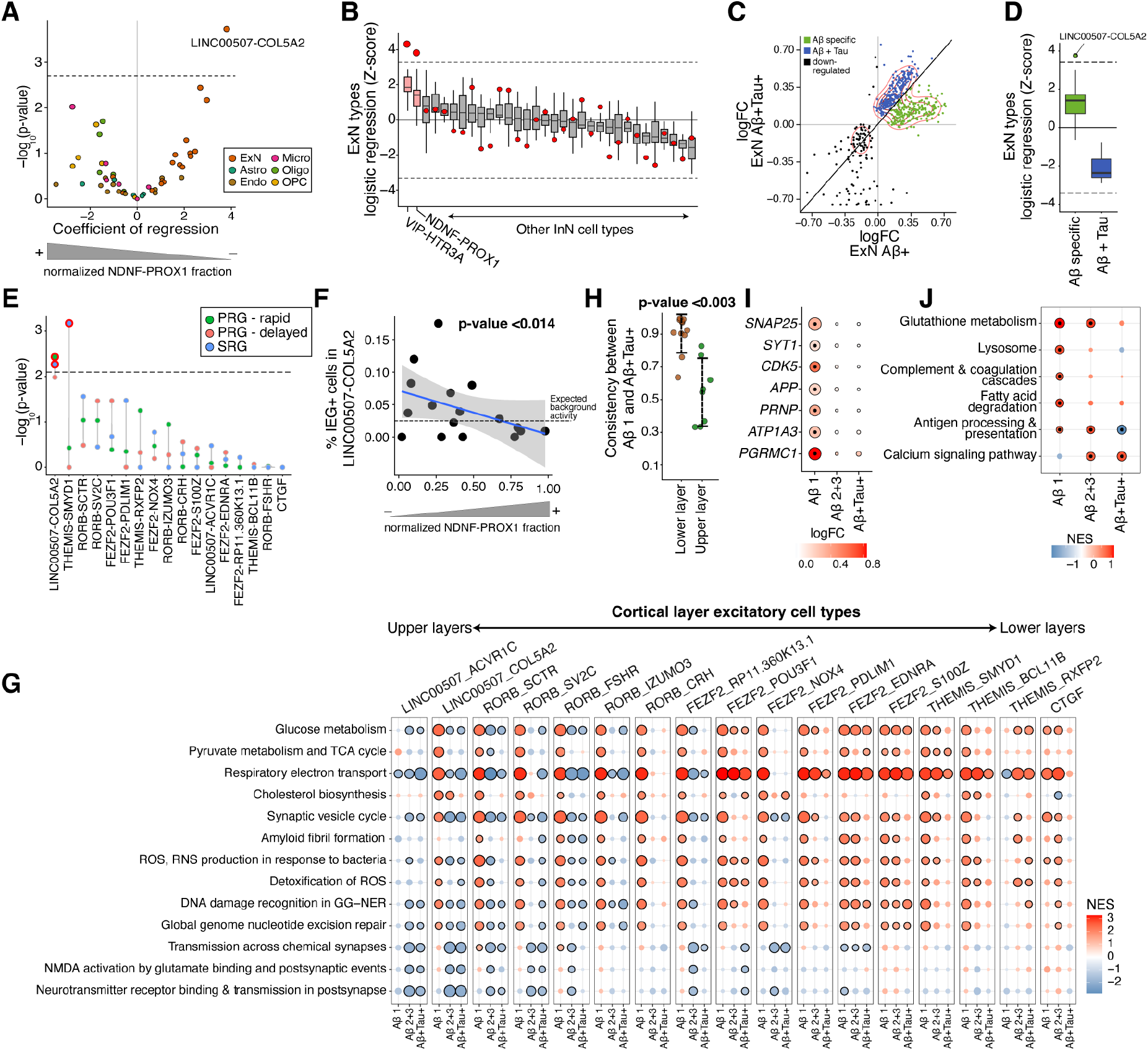
NDNF-PROX1 inhibitory neuron loss is associated with a hyperactivity signature in L2/3 excitatory neurons. ***A)** Logistic mixed-effect model regression of NDNF-PROX1 proportion versus cell type transcriptional signature in Aꞵ+ subjects. The dashed horizontal line represents the FDR threshold of 0.05. **B)** Associations (by logistic mixed-effect model) between the proportion of each inhibitory neuron type with each ExN type’s transcriptional signature in Aꞵ+ subjects. The red dots indicate the regression Z-score of the LINC00507-COL5A2 neurons with the corresponding inhibitory neuron cell type. The dashed line represents an FDR threshold of 0.05. Center line, median; box limits, upper and lower quartiles; whiskers, 1.5x interquartile range. See Figure S7H for more details. **C)** Scatter plot comparing the logFC in the ExNs of Aꞵ+ (x-axis) and Aꞵ+Tau+ samples (y-axis). Visualization is based on the union of top 300 protein-coding DE genes (sorted by jack-knifed p-value) in either group. **D)** Logistic mixed-effect model regression of NDNF-PROX1 proportion versus early-specific up-regulated DE genes (green dots in **C**) and up-regulated DE genes shared in both Aꞵ+ and Aꞵ+Tau+ samples (blue dots in **C**) for each ExN cell type. The dashed lines represent an FDR threshold of 0.05. Center line, median; box limits, upper and lower quartiles; whiskers, 1.5x interquartile range. **E)** Logistic mixed-effect model regression of NDNF-PROX1 cell fraction versus expression of neural activity signatures*^23^ in each ExN type in Aꞵ+***Continuation of*** Figure 3 ***legend.*** *samples (one-sided). The dashed line represents a one-sided FDR threshold of 0.05. PRG, primary response genes; SRG, secondary response genes. **F)** Scatter plot showing normalized NDNF-PROX1 fraction (x-axis) and the percent of LINC00507-COL5A2 ExNs with high expression of the core immediate early genes FOS, JUNB, ARC, NPAS4, ERG1, and ERG2 (y-axis, Methods) in Aꞵ+ subjects. A logistic mixed-effect model was used to calculate the p-value. **G)** GSEA of Reactome pathways on DE results from subjects with varying Aꞵ and tau burdens, across ExN types. Dots outlined in black denote significant terms (FDR-adjusted p-value < 0.05). Aꞵ+ individuals with Aꞵ burden scores of 2 and 3 are grouped together. **H)** Concordance of DE genes between different stages of AD pathology within excitatory neuron cell types. The LINC00507+ and RORB+ were selected as upper layer excitatory neurons and FEZF2+, CTGF+, and THEMIS+ populations as lower layer. **I)** ExN DE genes whose products are involved in synapse vesicle cycle and trafficking (SYT1, SNAP25, and CDK5), amyloid precursor protein (APP), and receptors of oligomeric Aꞵ (PRNP, ATP1A3, and PGRMC1) across different Aꞵ and tau burdens. The outlined dots represent DE genes with jack-knifed FDR-adjusted p-value < 0.01. **J)** GSEA of human KEGG gene sets using DE genes of WIF1+ homeostatic astrocytes across increasing Aꞵ and tau burden. Outlined dots represent significant terms (FDR-adjusted p-value < 0.1).*

We next sought to better understand the association of the ExN DE signature with Aꞵ plaque pathology. Comparison of ExN DE genes between Aꞵ+ and Aꞵ+Tau+ biopsies revealed a bimodal pattern among upregulated DE genes (Figure 3C), in which one DE gene set was evident solely in the early stage of pathology while another was present in both early- and late-stage samples. Given their differences in expression trajectory, we asked whether these two sets of DE genes were differentially correlated with NDNF-PROX1 inhibitory loss. Only the DE genes specifically found in response to early AD pathology, particularly within the LINC00507-COL5A2 population, correlated with NDNF-PROX1 proportional loss (Figure 3D).

The activity of layer 1 *NDNF-*expressing inhibitory neurons has been shown to play crucial roles in the integration of long-range inputs into cortex, particularly through gain modulation of whole cortical columns^24, 25^. We wondered if their loss may alter excitability of nearby L2/3 pyramidal cells. Indeed, we identified a significant association between NDNF-PROX1 loss and upregulation of neural activity response genes^23^ specifically within LINC00507-COL5A2 excitatory neurons in Aꞵ+ individuals (FDR-adjusted p-value < 0.01; Figure 3E). Furthermore, Aꞵ+ biopsy samples with a greater proportional loss of NDNF-PROX1 cells showed a higher percentage of LINC00507-COL5A2 cells expressing the core immediate early genes (*FOS, JUNB, ARC, NPAS4, ERG1*, and *ERG2)* that are induced after neuronal activity^26^ (p-value < 0.014; Figure 3F). Increased activity of excitatory neurons would be expected to affect their metabolism. Consistent with this, gene set enrichment analysis (GSEA) demonstrated increased expression of metabolism- and mitochondria-related gene sets (Methods) specifically in biopsies with the lowest level of Aꞵ plaque burden, further reinforcing the relevance of the hyperactivity phenotype to the early stages of AD pathology (Figure 3G). We also found an upregulation of gene sets indicating a cell-protective response to increased metabolism, including cholesterol biosynthesis, and responses to both reactive oxygen species (ROS) and DNA damage (Figure 3G). The enrichment of these terms was diminished in biopsies with higher burdens of Aꞵ and the presence of phosphorylated tau (Figure 3G), a pattern that was stronger in upper layer excitatory neurons. Consistently, comparing samples with lowest Aꞵ burden with Aꞵ+Tau+ demonstrated a significantly higher divergence of the DE patterns of upper layer neurons expressing *LINC00507* and *RORB* compared to the lower layer excitatory neurons expressing *FEZF2*, *THEMIS*, and *CTGF* (p-value < 0.003; Student’s t-test; Figure 3I), demonstrating the specificity of this response to upper layer cortical neurons at the early stages of Aꞵ plaque formation. Collectively, our results demonstrate NDNF-PROX1 inhibitory neuron loss is correlated with hyperactivity and preferential loss of layer 2/3 excitatory neurons in the prefrontal cortex with low Aꞵ plaque burden.

Hyperactivity of neurons can trigger homeostatic pre- and postsynaptic mechanisms^27^. In subjects with a low Aꞵ burden, we identified upregulation of *SNAP25, SYT1,* and *CDK5* in excitatory neurons, three genes whose products are involved in presynaptic vesicle release^28–30^ (Figures 3G and 3I). Increased activity of the presynaptic vesicle cycle can elevate Aꞵ production^31^. Congruently, we found upregulation of genes encoding for protein components involved in Aꞵ fibril formation, such as *APP* itself, only in the Aꞵ-low disease samples (Figures 3G and 3I). The oligomeric Aꞵ receptor genes *PRNP*, *ATP1A3*, and *PGRMC1*, whose protein products influence neuronal activity through the modulation of N-methyl-D-aspartate (NMDA) receptors^32^, were similarly upregulated in excitatory neurons at the early stages of AD pathology (Figure 3I). Homeostatic astrocytes also play critical roles in supporting synaptic function and coordinating antioxidant responses, especially in the context of neuronal hyperactivity^33, 34^. In our integrative analysis of astrocytes, we identified one *WIF1*+ type with low expression of *GFAP* and high expression of *EAAT1*, *EAAT2*, and *GSTP1* genes, which encode for critical components of glutamate/glutathione cycling (Figures S7I and S7J). The *WIF1*-expressing astrocytes showed enrichment of DE genes related to glutathione metabolism, lysosomal machinery, and fatty acid degradation specifically in subjects with low Aꞵ burden (Figure 3J), consistent with gene sets previously reported to be upregulated in the astrocytic response to hyperactive neurons^34^. Together, these results suggest that aberrant activity and metabolism of upper layer pyramidal cells perturb synapse homeostasis and astrocyte functioning in the brain.

### Expanded microglia populations with AD-specific alterations

Our integrative analysis across four AD-related cohorts indicated a mild expansion of *GPNMB*-expressing microglia population at early stages of AD pathology that further expands and becomes the strongest signal in samples with high histopathological burden (Figure 2A; Table S4). Human genetics and transcriptome studies have strongly implicated microglia in the AD pathogenic process^35–38^. A reactive population expressing *GPNMB* was also identified as enriched in an AD animal model near amyloid plaques^39^, but its connection to human *in vivo* microglial states–in AD, normal aging, and other diseases–remains debated. To more deeply explore microglial states in AD pathology, we leveraged our well-powered integrative analysis of 400,743 microglia profiles across human and mouse studies from diverse brain regions and biological conditions, including 59,624 high-quality microglia nuclei (median number of genes per nucleus = 2,384) from our biopsy cohort (Table S2). We more deeply sub-clustered the microglia profiles into a total of 13 microglial states (Figures 4A and 4B), including five homeostatic (HM) states, a chemokine-enriched state (CRM-CCL3), three reactive states expressing *GPNMB* (GPNMB-NACA, LPL-CD83, and GPNMB-EYA2), an interferon gene-enriched state (IRM-IFIT2), and a proliferative (Prolif) state (Figures 4A and 4B; Table S3). Comparing the microglia populations, we observed the main microglial markers, including *SLC2A5*, *CX3CR1*, *CSF1R*, *P2RY12* were downregulated in the three *GPNMB*-expressing populations relative to the homeostatic microglial cells, but were still expressed at higher levels than macrophages and myeloid cells (Figure 4A). One of the smaller homeostatic microglia populations, which we designated HM-2, also exhibited high expression of genes associated with technical dissociation artifacts, including *FOS* and *JUNB*^19^. All microglia states were well represented across datasets, biological conditions, and sequencing platforms. Moreover, we observed that markers of microglia states correlated strongly across different human brain regions, which is consistent with previous reports^40, 41^ (Figures S8A-S8D).

**Figure 4.**
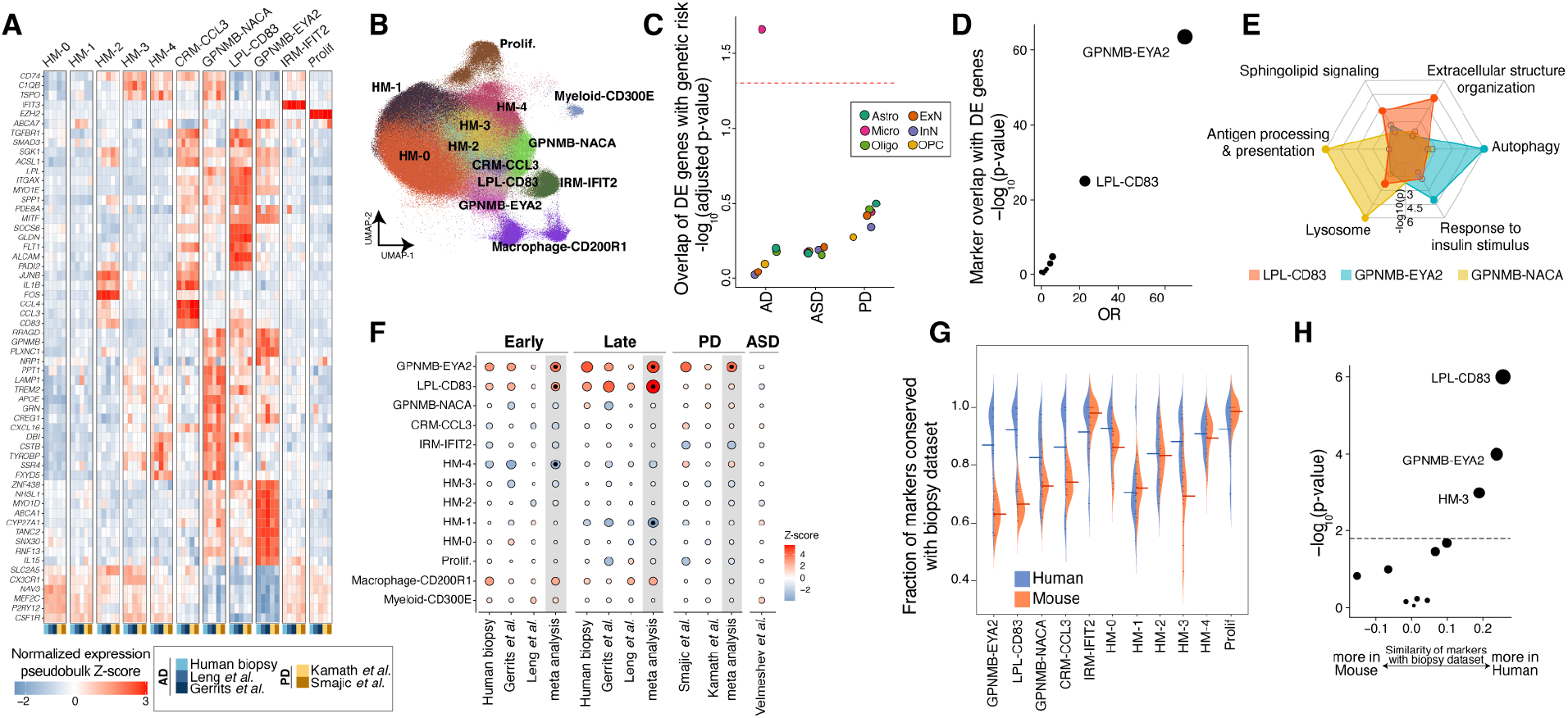
Precise molecular definitions of microglial states activated in early and late AD. ***A)** Expression of select marker genes (Methods) across human neurodegeneration datasets in the microglia integrative analysis. The expression values represent pseudobulk expression of each marker in each cell state and dataset. **B)** Uniform manifold approximation and projection (UMAP) representation of microglia profiles from integrative analysis, colored by the 13 identified states. **C)** Dot plot of -log*^10^*-transformed p-values for MAGMA enrichment analysis*^42^ *(y-axis) of AD, PD, or ASD genetic risk in the up-regulated DE genes of each cell class. Dots are colored by cell class membership. Dashed line represents an FDR threshold of 0.05. **D)** Dot plot of -log_10_-transformed p-values for a Fisher’s exact test assessing the overlap between microglial DE genes with markers of each of the 13 microglial states (Methods). **E)** Radar plot representation of enriched gene sets in markers of the three GPNMB-LPL states. The marker analysis was conducted by comparing the three cell states against each other. See **Table S6** for more details. **F)** Association of proportion of each microglial state with early and late AD pathology, as well as PD and ASD. In meta-analysis columns, black dots represent microglia states with significant changes in cell state proportions (FDR-adjusted p-value < 0.05). The scale of points is based on the absolute Z-score values. **G)** Distribution of the fraction of markers shared between the biopsy cohort and each other dataset (y-axis), in each microglial state (x-axis). Datasets are stratified by species. Mean values are denoted with a line. Only genes expressed in more than 1% of cells were considered in the analysis of each dataset. **H)** Statistical comparison of the differences in (**G**) by Student’s t-test. The dashed line represents an FDR threshold of 0.05*.

Next, we examined how each of these 13 microglial states was affected by the presence of Aꞵ. Differential expression analysis across all microglia in our cohort identified a pattern that was highly similar in each of the 13 states (Figures S8E and S8F), suggesting that all microglia states respond to Aꞵ accumulation in a similar manner. This transcriptional pattern was also highly consistent across postmortem cohorts (Figure S8G). The DE signature showed upregulation of genes whose protein products are involved in microglia neuroinflammatory responses, including phagocytosis (*COLEC12*), antigen presentation (*CD74* and *HLA* genes), lipoprotein metabolism and biosynthesis (*APOE*, *OLR1*, *ATG7*), fatty acid metabolism (*ACSL1*), autophagy (*ATG7*, *ATG16L2*), and lysosomal function (*ASAH1, NPC2, SLC11A1, PSAP*) (Table S5). Underscoring the pathological relevance of this common microglial DE signature, we found that it was significantly (FDR-adjusted p-value < 0.05) enriched for the expression of genes implicated in AD by common variant case-control studies^35, 36^, including: *APOE*, *MS4A6A/4A*, *TREM2*, and *INPP5D* (Figure 4C, Methods). In addition, intersection of this DE signature with marker genes for each of the 13 states showed highly significant overlap with markers of GPNMB-EYA2 and LPL-CD83 microglia (FDR-adjusted p-value < 0.001; Fisher’s exact test; Figure 4D), indicating a transcriptional transition across microglia cells towards a state more resembling the GPNMB-EYA2 and LPL-CD83 populations. These results indicate that microglia cells collectively transition towards a transcriptional state with high expression of AD risk genes and neuroinflammatory-related processes as AD pathological burden increases in human brains.

**Figure 5.**
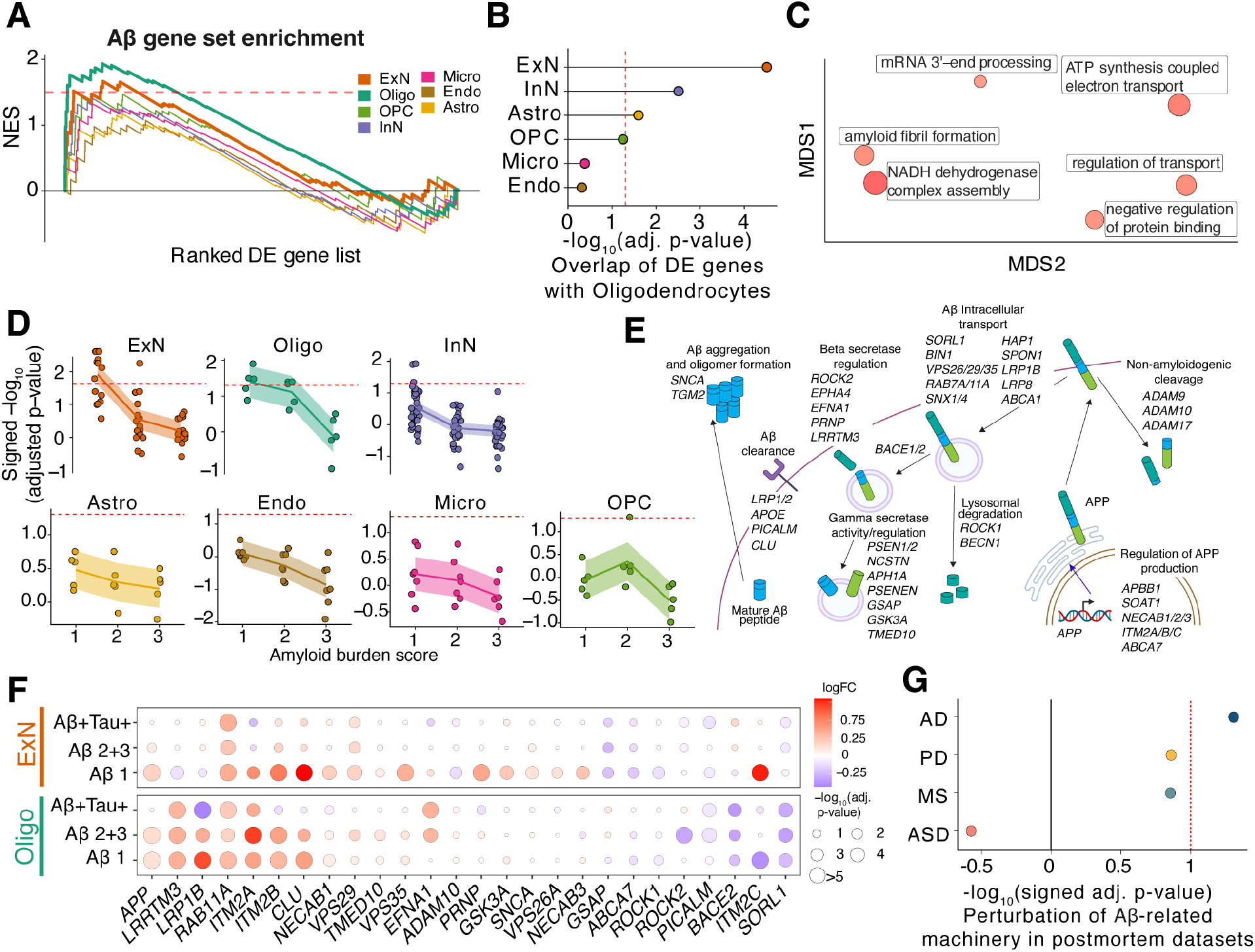
Cell-type-specific dysregulation of amyloid formation in the human frontal cortex. ***A)*** *GSEA trace plot of amyloid-associated gene set ordered by their signed p-value from DE analysis across the seven cell classes. The x-axis shows the rank order of the DE genes in corresponding cell classes; the y-axis is the normalized enrichment scores (NES) from GSEA. Bold lines indicate GSEA traces for significant cell classes, oligodendrocytes and excitatory neurons. The dashed line indicates NES score corresponding to FDR threshold of 0.05. **B)** Dot plot of -log_10_-transformed FDR-adjusted p-values of GSEA results of the top 300 upregulated protein-coding genes (sorted by their jack-knifed p-values) from each cell class against an ordered list of DE genes in oligodendrocytes. Dotted red line indicates significance at FDR threshold of 0.05. **C)** Multidimensional scaling (MDS) low-dimension embedding of gene ontology terms significantly enriched in intersect of DE genes between oligodendrocytes and excitatory neurons from REVIGO*^46^ *(see Methods). Size of dots indicate significance values. **D)** GSEA of amyloid gene set against cell type level DE genes across increasing levels of Aꞵ and tau burden. Cell types are grouped based on their major cell class annotations. The dashed line represents a significance threshold of FDR-adjusted p-value < 0.05. **E)** Schematic of regulation of Aꞵ formation, intracellular transport, and degradation/clearance pathways, showing the substituent in each pathway genes that together comprise the amyloid gene set (**Table S7**). **F)** Excitatory neuron and oligodendrocyte DE results across increasing levels of Aꞵ and tau burden for genes found by the leading edge analysis in **A**. The size of each dot is scaled by p-values and the color of each dot denotes the logFC. **G)** Signed -log_10_-transformed p-values from GSEA results* for the amyloid gene set on *Oligo DE genes from postmortem AD, PD, MS, and ASD cohorts*.

We focused particularly on the three reactive *GPNMB+* states, given their disease relevance. All three states highly expressed genes related to microglial reactivity, including *APOE, ITGAX, MITF,* and *SGK1* (Figure 4A). However, comparative marker analysis between the three states revealed substantial differences. The GPNMB-NACA population preferentially expressed genes involved in antigen processing and presentation, as well as lysosomal and phagosomal function relative to the other two states (Figure 4E). By contrast, the GPNMB-EYA2 microglia preferentially expressed genes involved in autophagy (e.g., *IGF1R*, *ATG7*, and *ATG16L2*) and response to insulin (e.g., *MYO5A*, *IGF1R*, and *PPARG*) (Figure 4E). This cell state also expressed *IL15*, a key modulator of the nervous system inflammatory response^43^ (Figure 4A). The LPL-CD83 microglia expressed genes, including *TGFBR1* and *SMAD3,* which encode for key proteins in TGF-ꞵ signaling (Figure 4A), and showed enrichment of genes involved in extracellular structure organization, response to cytokines, focal adhesion, and actin cytoskeleton (Figure 4E). Both IL-15 and TGF-ꞵ also mediate neuroinflammatory cross-talk between astrocytes and microglia^44, 45^. Supporting this notion, we found a strong positive correlation between the expression of *GFAP* in astrocytes and the expansion of GPNMB-EYA2 and LPL-CD83 microglia states in our cohort (Figure S8H). Together, we find transcriptional heterogeneity among reactive microglia cells in human brains that points to specialized functional roles in responding to cues from their surrounding microenvironment.

We next conducted a meta-analysis to ask which of these microglial states is specifically enriched in AD, and how these states relate to those found in other neurodegenerative diseases and disease models. Across the three AD-related datasets with sufficient numbers of microglia to power proportional testing, we identified an expansion of the LPL-CD83 and GPNMB-EYA2 states in both early and late stages of AD histopathology (FDR-adjusted p-value < 0.05; Figure 4F). Interestingly, the GPMNB-EYA2 state was also enriched in a meta-analysis of two PD datasets (FDR-adjusted p-value < 0.002; Figure 4F), while LPL-CD83 was exclusively expanded across the AD datasets. Neither *GPNMB*-expressing microglia population was expanded in individuals with ASD, underscoring the specific role of these microglia in neurodegenerative diseases. In contrast to the human datasets, only the GPNMB-NACA state was consistently expanded in AD mouse models (Figure S8I). This state was also increased in several other mouse datasets, including a model of amyotrophic lateral sclerosis, in both juvenile and aged mice, and in response to demyelinating injury (Figure S8I). To better understand the underlying factors contributing to this apparent divergence in microglia response, we performed a systematic marker analysis of the 13 microglia states across the human and mouse datasets that are included in our integrative analysis. First, given the superior size, quality, and coverage of our human biopsy dataset, we used it as the base to reliably identify markers of each state. We next examined the concordance of marker genes in each of the remaining datasets. As expected, we found that microglia states are highly consistent across human datasets. Although human microglia states were less preserved in the mouse datasets in overall, preservation was notably lower for the mouse LPL-CD83 and GPNMB-EYA2 states (Figures 4G and 4H), suggesting these transcriptional states are less well recapitulated by laboratory mice. Collectively, our results demonstrate shared and AD-specific microglia responses to disease in the human brain, and selective divergence of the most disease-relevant states in mouse models.

### Amyloidogenic cell populations in human frontal cortex

The production of amyloid in the brain has largely been assumed to be only in neurons but has been challenging to directly study in human tissue. We leveraged our high-quality surgical biopsy dataset to assess amyloidogenicity in each cell type using transcriptional signatures as a proxy. We took an unbiased approach, assessing the enrichment of a set of 49 genes known to regulate Aꞵ production and secretion in each cell type (Table S7). Interestingly, GSEA against an ordered list of DE genes for each of the seven cell classes identified not only excitatory neurons but also oligodendrocytes as having significant, positive enrichment for the amyloid gene set (FDR-adjusted p-value < 0.05, Figure 5A, Methods). The enrichment in these two cell classes was robust to the statistic used to order genes (Figure S9A). A leading-edge analysis identified specific genes that were highly up-regulated (including *APP*, *LRRTM3*, and *ITM2B*) and down-regulated (such as *BACE2, SORL1*, and *PICALM*) in biopsy samples with Aꞵ plaques in our cohort (Figure S9B).

The unexpected enrichment of amyloid-related genes in oligodendrocytes prompted us to investigate whether they share a common DE gene signature with excitatory neurons. Assessment of the degree of overlap between DE genes from oligodendrocytes and other cell classes revealed that the excitatory neuron DE genes had the most significant degree of overlap (FDR-adjusted p-value < 0.05; Figure 5B), which was consistent across a wide range of gene set sizes tested (Figure S9C). The high overlap of DE genes between oligodendrocytes and excitatory neurons is in contrast with the overall expression of genes in oligodendrocytes, which overlaps most with OPC populations (Figure S9D), indicating a selective dysregulation of shared processes in oligodendrocytes and excitatory neurons. A gene ontology analysis of the intersecting co-regulated genes identified enrichment for multiple terms, including those related to amyloid fibril formation (Figure 5C), further suggesting a similar, shared Aꞵ-related response. Remarkably, this signature was most prominent in the samples with lowest Aꞵ burden for both excitatory neurons and oligodendrocytes (FDR-adjusted p-value < 0.05, Figures 5D, S9E and S9F). The DE genes that made up the leading edge from GSEA were involved in multiple aspects of amyloid processing, including regulation of *APP* transcription, beta-secretase regulation, and degradation/clearance pathways for Aꞵ peptides (Figure 5E). These changes included the downregulation of genes known to decrease the production or else help clear Aꞵ peptides, such as *SORL1*, *BACE2*, and *PICALM*, as well as the upregulation of genes involved in amyloid formation like *RAB11A, LRRTM3,* and *APP* itself (Figure 5F). Crucially, the Aꞵ gene set was consistently enriched across oligodendrocytes in a meta-analysis of postmortem cohorts with low AD histopathology^3, 4^ (Figure S9G), and not in DE genes from other disease states, including PD^47, 48^, ASD^49^, and MS^50^ (Figure 5G), reinforcing its robust and specific association with AD across cohorts.

To experimentally assess the relative Aꞵ-forming potential of these two cell populations, we differentiated the H1 embryonic stem cell (ESC) line into mature oligodendrocytes (iOligos) and excitatory neurons (iExNs) (Figure 6A; Methods). In our iExN culture, we found nearly all cells expressed major excitatory neuron markers, as measured by single-cell RNA-sequencing (Figures S10A-S10C). Single-cell analysis (Figure 6B) of our iOligo cultures showed robust expression of numerous genes known to play roles in myelin function such as *MBP, PLP1*, and *CNP*, as well as transcription factors important for oligodendrocyte differentiation and maturation such as *SOX10* and *NKX2-2* (Figures 6C and S10D-S10F). These genes were not expressed at high levels in our excitatory neuron cultures, which instead were marked by canonical neuronal marker genes such as *RBFOX3, SLC17A7*, and *TUBB3* (Figure 6C). Importantly, both cultures expressed appreciable levels of all the necessary machinery to produce amyloid beta protein (Figure 6D). Further, immunohistochemistry of key proteins defining the oligodendrocyte lineage, such as *MBP* and *O4*, showed a linear, significant increase upon induction of *SOX10* (Figure 6E, Figure S10I, p-value < 0.05, linear mixed-effect model), while markers of other cell types were not significantly associated (Figures S10G and S10H).

**Figure 6.**
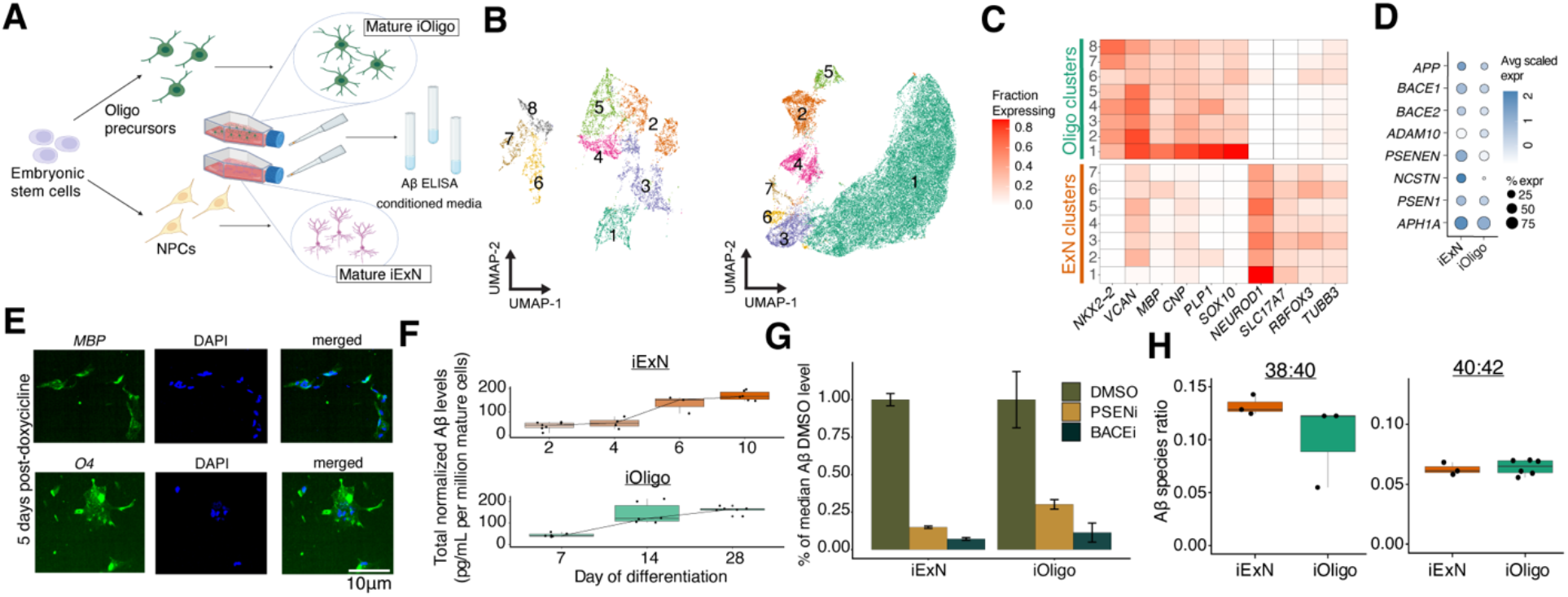
Quantitation of Aꞵ production by human mature oligodendrocytes and excitatory neurons. ***A)** Schematic of differentiation of ESCs and ELISA-based quantification of Aꞵ from conditioned media. **B)** Two-dimension UMAP embeddings of single-cell expression profiling for ESC-derived iOligo (left) and iExN (right) cultures. **C)** Expression of key marker genes in ESC-derived iOligo and iExN cultures. **D)** Dot plot depicting scaled expression of essential Aꞵ machinery in ESC-derived cultures of iOligos and iExNs. **E)** Representative images of immunofluorescence stains of O4 and MBP in ESC-derived iOligo cultures five days after doxycycline addition. **F)** Normalized Aꞵ protein abundance for ESC-derived iExNs (top) and iOligos (bottom) across days of differentiation. Center line, median; box limits, upper and lower quartiles; whiskers, 1.5x interquartile range; points, outliers. **G)** Fractional abundance of Aꞵ protein levels relative to median Aꞵ protein levels in DMSO condition for PSEN inhibitor-treated and BACE inhibitor-treated conditioned media samples for ESC-derived iOligos and iExNs. Error bars indicate one standard deviation above and below the mean value. **H)** Ratio of Aꞵ-38 to Aꞵ-40 and Aꞵ-40 to Aꞵ-42 species from conditioned media obtained from ESC-derived cultures of iExNs (left) and iOligos (right). Center line, median; box limits, upper and lower quartiles; whiskers, 1.5x interquartile range; points, outliers.*

An ELISA-based quantification of amyloid-beta peptides from our iOligo cultures (Methods) showed a linear, three-fold increase in Aꞵ upon *SOX10* induction (Figure 6F). We observed a similar fold increase after induction of differentiation in the iExN culture (Figure 6F). Indeed, the total abundance of Aꞵ was not significantly different (p = 0.497, Student’s t-test, Figure 6F) between iExN and iOligo at their respective differentiation endpoints, suggesting a similar intrinsic cell-autonomous capacity to produce amyloid beta. As predicted with our analysis of single-nucleus transcriptome data, we could not detect any Aꞵ in media taken from ESC-derived microglia cultures, with raw values similar to those found in an unconditioned media (Figure S10J). Further supporting the functionality of Aꞵ production and processing machinery in both populations, treatment of both iOligo and iExN cultures with a beta-secretase (BACE) inhibitor or gamma-secretase inhibitor caused a 10-fold reduction in total Aꞵ protein levels in both cell types (Figure 6G). Finally, we sought to determine whether the species composition of amyloid peptide production was significantly different between cell types given the well-established higher aggregation and amyloid formation potential of longer species. We found that Aꞵ species ratios were not significantly different (p-value = 0.64, Student’s t-test, Figure 6H) between iOligos and iExNs, suggesting that oligodendrocyte-derived Aꞵ peptides could contribute to AD-related amyloidosis in similar ways to excitatory neurons.

## Discussion

Therapeutic trials of AD have made increasingly clear the importance of early intervention into the disease^2^, but identifying the cellular states occurring in human tissue at early disease stages has been challenging. Here, we leveraged a unique cohort of fresh human brain biopsy tissue to identify cellular perturbations–which we collectively refer to as the Early Cortical Amyloid Response (ECAR)--that are specifically present in tissue at the earliest stages of AD pathology. One prominent ECAR component was the identification of a hyperactive, hypermetabolic signature within excitatory neurons. This signature was associated with an astrocytic upregulation of glutathione metabolism and fatty acid degradation, suggesting dysregulation of synapse homeostasis in response to aberrant neuronal activity^33, 34^. Furthermore, this upper layer hyperactivity phenotype was tightly coupled with the loss of a specific *NDNF+* layer 1 interneuron population very early in disease progression. *NDNF*-expressing interneurons are most active in states of arousal^25^, and their activation is positively correlated with associative learning^51^, suggesting that their loss may directly affect memory formation. In addition, a recent study of *NDNF*-expressing neurons in the hippocampus found that their potentiation led to an inhibitory shift at excitatory synapses between entorhinal cortical projections and CA1 neuronal dendrites^52^. The loss of this cell type could thus help seed foci of aberrant excitation, further aggravating the acute effects of Aꞵ accumulation in the tissue. Hyperactivity has been observed in animal AD models that either overexpress APP or are exposed to Aꞵ-containing extracts from AD patients^32, 33, 53, 54^. Our work in human tissue finds that hyperactivity is a prelude to subsequent neuronal loss, and postulates a mechanism–loss of a specific inhibitory neuron population–that contributes to its onset.

The second ECAR component is the expansion of two activated microglial states, one of which (GPNMB-EYA2) is shared between AD and PD, and the other (LPL-CD83) that is expanded only in AD. One means by which microglia protect against neurodegeneration is through the autophagy-mediated clearance of Aꞵ^55^ and α-synuclein^56^, a convergence that could explain the expansion of the GPNMB-EYA2 population–enriched for autophagy-related genes–in both AD and PD. The LPL-CD83 population–whose expansion is AD-specific–shows high expression of TGF-ꞵ signaling components, including *TGFBR1* and *SMAD3*, which both promote Aꞵ clearance by microglia^57^ and mediate tissue repair^58^. According to our integrative analysis, neither of these cell states was expanded in the examined AD mouse models, and the states themselves showed more molecular divergence between species than other microglial states, arguing for the importance of human samples when studying these highly disease-relevant cells. Interestingly, exposure of human ESC-derived microglia to diverse brain-related challenges was recently shown to induce *in vitro* cellular states that transcriptionally resemble our *GPNMB+* states (e.g., high expression of *GLDN*, *CD83*, *PPARG*, and *MYO1E*)^59^. It will be important to more deeply characterize these states genomically, and to study their functional properties, such as capacity for phagocytosis, synaptic engulfment, and neuroinflammatory potential.

The last ECAR component we identified was a shared signature, in both oligodendrocytes and excitatory neurons, of differentially regulated genes associated with Aꞵ production. This signature peaked especially in the lowest-stage amyloid burden samples, suggesting a declining rate of amyloid production with the progression of the disease, consistent with rates of amyloid accumulation determined from pre-clinical non-invasive measurements made in patients with dominantly-inherited AD^60^. Because our signature derives from measurements made from human biopsy tissue, it provided us with a unique opportunity to uncover the molecular mechanisms underlying excess production and accumulation of Aꞵ in early stages of AD pathology in the human brain. Upregulated genes in this signature encoded for pro-amyloidogenic factors such as *LRRTM3,* and *RAB11A,* as well as *APP* itself, while *ITM2B*, *SORL1*, and *BACE2* were downregulated. The dysregulation of these specific genes within human diseased tissue nominates them as especially promising targets for therapeutic intervention into early amyloid beta accumulation.

Our analyses establish oligodendrocytes as an amyloid-producing cell type in AD. Prior work in animal cells and models have suggested that other cell types, beyond excitatory neurons, could be sources of amyloid^61, 62^. Our work–supported by both analyses of human tissue and human ESC-derived cultures–suggests that in humans, Aꞵ production is primarily in oligodendrocytes and excitatory neurons. Neuropathological studies have postulated an inverse relationship between myelination and AD pathology^63^, prompting hypotheses that myelin breakdown may play a causal role in the disease^64^. Additionally, white matter regions are some of the first to exhibit a high burden of oligomeric Aꞵ^65^. These studies, coupled with our results, underscore the relevance of the interface between neuronal axons and oligodendrocytic bodies to AD pathogenesis.

Although our findings were enabled by the exceptional data quality obtained from a rare cohort of freshly-frozen brain biopsies, we utilized large-scale integrative analysis of many published datasets to corroborate many of our findings, and to assess their specificity for AD. Our integrative analysis illustrates that cell type identities are more resilient to peri- and post-mortem effects compared to expression patterns of individual genes, and can be accurately recovered by anchoring to high-quality datasets. From this work, we conclude that single-cell brain datasets are generally of sufficient consistency and quality that it is possible to conduct cumulative, highly informative meta-analyses. Such analyses will not only ensure the consistency of biological findings across multiple cohorts, but will also enable comparative analyses–as we performed here across many diseases and models with microglial expansion phenotypes–to assess the specificity of a state for a particular disease. To further facilitate this endeavor, we have established a web-based resource (available at https://braincelldata.org/resource) where individual scientists can explore our integrative analysis–which covers most published single-cell studies of the brain– to formulate and test mechanistic hypotheses. In addition, we have generalized the process of data integration to enable scientists to seamlessly integrate their own new datasets into our analysis, providing a common language for understanding cell-type-specific changes in different cortical diseases. We expect that the continued accrual of data from more donors, regions, species, and related conditions will provide additional crucial insights into the pathogenic process of AD and other diseases of the brain.

## Acknowledgments

We would like to thank Henrik Zetterberg, Nader Morshed, and Michael-John Dolan for helpful discussions. We are also grateful for the help of Marita Parviainen and Tiina Laaksonen with patient management and cognitive testing, and Andrea Jiang with performing the amyloid beta ELISA assays. This work was supported by the Alzheimer’s Association and the International Neuroimmune Consortium (to B.S. and E.Z.M.), the Pew Biomedical Scholars Program (to E.Z.M.), NIH NIGMS grants T32GM007753/T32GM144273 (to T.K.), NIH NIA grant F30AG069446 (to T.K.), and the Kuopio University Hospital VTR fund (grant 5252614 to V.L.); Academy of Finland (grants 339763, 334801 to TM; 338182, 328287 to MH); the Sigrid Juselius Foundation; the Strategic Neuroscience Funding of the University of Eastern Finland, the Finnish Cultural Foundation, and the North Savo Regional Fund and by the European Union (ERC, HUMANE, 101043584 to TM). Views and opinions expressed are however those of the author(s) only and do not necessarily reflect those of the European Union or the European Research Council Executive Agency. Neither the European Union nor the granting authority can be held responsible for them.

## Author contributions

V.G. and T.K. performed all analyses, with help from A.J. and S.M. N.N. generated the snRNA-seq data, with help from T.K., C.V. and A.A. V.L. conceived, designed and oversaw the construction of the biopsy cohort, with help from M.H. and T.M. T.K., B.R., M.Th, M.Te, and R.N. performed the differentiated stem cell experiments. T.R. pathologically staged the biopsies and S.H. collected the CSF from patients. B.S., V.L., and E.Z.M. conceived of the study. V.G., T.K., and E.Z.M. wrote the paper, with contributions from all authors.

## Competing interests

The authors declare no competing interests.

## Data availability

All generated snRNA-seq data and the results of our integrative analysis of 28 single cell/nuclei studies are publicly available at: https://braincelldata.org/resource. This includes sample annotations related to the dataset source (36 datasets across 28 studies), cell identifiers (e.g., cell barcodes), quality metrics, and cell type annotations from integrative analysis. We have also included functionality to perform several analyses in a fast and efficient way, including: examination of the integration solutions, performing marker analysis across all of the datasets, and exploring differentially expressed genes.

## STAR Methods

### Procurement of frontal cortex brain biopsies

Patients presenting to a clinic at the Kuopio University Hospital were evaluated for adult hydrocephalus with NPH symptoms: 47 with idiopathic normal pressure hydrocephalus, 3 with previously unrecognized congenital hydrocephalus and 2 with acquired hydrocephalus. Patients were consented for retrieval of brain biopsies during ventriculoperitoneal shunt placement for treatment of their symptomatic adult hydrocephalus. Biopsies were taken at the site where the shunt would penetrate the brain. Three cylindrical biopsies were taken approximately 2mm in diameter and 3-10mm in length using a disposable Temno Evolution TT146 (Merit Medical Systems) biopsy tool. The insertion point of the catheter was approximately 3 cm from the midline and anterior to the coronal suture^66^. Biopsies were immediately frozen with liquid nitrogen and stored at -80°C. One biopsy was sent for histopathological staining using the 6F3D and AT8 antibodies and evaluated by a neuropathologist for presence of Aꞵ plaques and tau tangles via light microscopy^67^. Biopsy Aꞵ plaques burden was further assessed semiquantitatively by a neuropathologist (T.R.) under light microscopy and assigned to mild (1), moderate (2), or severe amyloid burden (3) as described previously^68^. Our initial cohort included 58 individuals. We excluded one individual with tau-only pathology and another patient with a history of psychosis. We excluded four additional individuals (2 Aꞵ-free and 2 Aꞵ+) that, upon microscopic inspection of Nissl stained cryosections (see the biopsy tissue quality scoring methods), displayed decidedly poor tissue quality and a very high ratio of white matter to cortical matter. All of these excluded biopsies were more than 85% white matter tracts with the diminished cortical regions showing dysmorphic neuronal profiles. The biopsy procedure was approved by the Research Ethics Committee of the Northern Savo Hospital District (decision No. 276/13.02.00/2016).

### Neuroanatomical localization of biopsy site

The stereotactic position (distances in millimeters) was measured from anatomically linked planes (transverse, sagittal, coronal) in a multiplanar reconstruction (MPR) produced from the postoperative CT/MRI DICOM image. After planar alignment (transverse and sagittal planes to the midline, and coronal plane in a 90-degree angle to the planum sphenoidale), the biopsy location was determined to be at proximal catheter’s cortex entry site at the catheter’s midline. Following distances were measured: In the transverse plane from the midline to the biopsy location (x). In the sagittal plane from the frontal bone’s internal cortex to the biopsy location’s coronal axis (at 90-degree angle) (y). In the sagittal plane, distance from the planum sphenoidale to the biopsy location’s transverse axis (at 90-degree angle) (z). EBRAINS Siibra-explorer was used to map and visualize each biopsy position^69–71^.

### Biopsy tissue quality scoring

To ascertain tissue quality measurements (range from 1-10), we performed Nissl staining followed by semi-quantitative scoring of each biopsy slide image. For Nissl staining, briefly, fresh frozen tissue was thermally equilibrated to -20°C in a cryostat (Leica CM3050S) for 20 minutes. Tissue was mounted onto a cryostat chuck with Optimal Cutting Temperature compound (O.C.T. compound), aligned at a 5° cutting angle, and sectioned at 10 μm in thickness per tissue slice. Using a Superfrost plus slide that has been pre-cooled to -20°C, the tissue section was collected by carefully placing it and gently flattening it with the brush on top of the slide. Subsequently, slides were stained with 0.1% Cresyl Violet acetate in DiH_2_O, destained with ethanol, and 100% Xylene for 5 minutes. Slides were mounted by adding 2-3 drops of Permount around the tissue and coverslipped. Images were subsequently collected using a Keyence BZ-X810 series All-in-one Fluorescence microscope. With BZ-X800 viewer software, each stained slide was imaged on the Brightfield/Phase contrast channel using a 20X objective. The stained region of interest was selected by specifying the XY positions of the tissue outer edges and adjusting the Z-stack function to auto-focus prior to each image capture. The stitching of captured image series was made with BZ-X800 analyzer software. The images are exported as Big TIFF files and edited for cropping, contrast, and brightness with Photoshop software.

### Measurement of iNPH grading scale

Severity of iNPH related symptoms was evaluated using the iNPH grading scale (iNPHGS)^72^, a clinician-rated scale that aims to assess the hallmark triad of the disease. Inversely-correlated with the severity of the disease, the 12-point iNPHGS has been shown to be clinically meaningful down to a single point^72^.

### Generation of single-nuclei suspensions from frozen brain biopsies

Fresh-frozen brain biopsy tissue was cryosectioned at -15 to -20°C into 60-micron sections. Following microdissection, samples were placed on dry ice until nuclei isolation. To each cryosectioned sample, 1 mL of Extraction Buffer (ExB) was added into a 1.5-mL Eppendorf tube. Samples were briefly triturated before being placed in a six-well plate. Samples were then triturated 20 times with the ExB, every 2 minutes, until no large chunks of tissue were observed in each well. After the last trituration, samples were diluted with 45-50mL of wash buffer in a 50-mL Falcon tube, and then split into four 13-15 mL solutions in 50mL Falcon tubes. The diluted samples were then spun at 500g for 10 minutes at 4°C (pre-cooled) in a swing bucket benchtop centrifuge.

After centrifugation, a visible nuclei pellet was observed. Samples were then removed very gently from the centrifuge and placed in an ice bucket. The supernatant was aspirated until there was barely any liquid observed on top of the pellet (50-100µL of liquid left). To aspirate without disturbing the pellet, a serological pipette was first used till about 1mL was remaining, followed by serial aspiration with a P2000 and P200 pipette.

The pellets were then resuspended in 250µL of wash buffer (WB), mixed thoroughly by trituration and placed in an Eppendorf 1.5-mL tube.

### Single-nucleus and single-cell RNA-sequencing and read pre-processing

For all single-nuclei experiments, the 10X Genomics (v3) kit was used according to the manufacturer’s protocol recommendations. Library preparation was performed according to the manufacturer’s recommendation. Libraries were pooled and sequenced on either a NovaSeq S2 or S4.

Sequencing reads from human brain biopsy experiments were demultiplexed and aligned to the hg19 reference using DropSeqTools (https://github.com/broadinstitute/Drop-seq) with the default settings. To reduce background noise from ambient RNA and potential UMI barcode swaps, we used Cellbender remove-background v2^73^ with the default applied settings to all libraries. The Cellbender-corrected reads were used for downstream variable gene selection, dimensionality reduction, clustering, and differential expression. Cellbender was also used to distinguish cells from empty droplets.

### Initial clustering of the biopsy cohort

Pre-processed Cellbender-corrected digital expression matrices were loaded into R per library as a digital gene expression matrix. All matrices were combined per individual and an initial variable gene selection was performed. A low-dimensional embedding was generated via rliger v1.0 at a k = 45 and lambda = 5. Following integrative non-negative matrix factorization^18^, a shared nearest neighbors graph was generated and individual nuclei profiles were clustered according to the SLM (smart local moving) algorithm to identify broad cell classes. We used a recent large-scale survey of postmortem human brain^21^ to identify cell class markers and merged each cluster into one of eight cell classes (excitatory neurons, inhibitory neurons, astrocytes, microglia/macrophages, oligodendrocytes, oligodendrocyte precursor cells, endothelial cells/pericytes, and peripheral blood mononuclear cells (PBMCs)). PBMCs were excluded from downstream analysis.

For each cell class, individual nuclei were subsetted and the above clustering process was repeated to identify individual cell types. Marker genes were identified for neuronal populations based on a large-scale survey of neurons in the human neocortex^21^. Further, a recent survey of microglia/macrophage in the murine brain was used to identify cell type markers for microglia and macrophages^74^. For other non-neuronal types, we performed the Wilcoxon rank-sum test on SLM-defined cell clusters to find markers and thereby determine cell type annotations. We removed doublets identified as clusters that expressed markers of more than one cell class population. We also removed clusters whose markers contained high numbers of mitochondrial genes or heat shock related proteins.

### Integrative analysis of the biopsy dataset with postmortem studies

We collected and uniformly processed all publicly available metadata on each dataset including the donor information (e.g., age, sex, diagnosis), sample information (e.g., brain region, sequencing protocol, batch structure), cell type identities, and quality metrics. All gene identifiers were mapped to Ensembl gene id. For mouse datasets, we further mapped Ensembl gene ids to their human orthologs^75^. However, we did retain non-orthologous mouse genes for normalization. We calculated the following quality metrics for every cell in each dataset: number of unique genes (nGene) and total unique molecular identifier (nUMI), percentage of mitochondrial genes (MT%), percentage of ribosomal genes (Ribo%), percentage of non-coding lncRNAs (lncRNA%), and percentage of dissociation-related artifact genes^20^. We used nGene and MT% quality metrics as our initial criteria to select cells for our integrative analyses and used the other quality metrics to identify and remove low quality cell clusters from the integrative analysis results. We retained cells with nGene >500 and MT% <5. For microglia cells, we used nGene >200 for two studies^4, 39^ to compensate for the lower number of unique genes compared to other cell types. We further used the nGene >200 threshold for all cell classes in Mathys *et al.* dataset^3^. Finally, we removed donors with less than 50 cells within each cell class. Our integrative analyses across the seven cell classes included a total of 2,406,980 high quality cell profiles across 36 datasets from 28 studies on humans and mice. Distributions of the quality metrics are included in Figures S3 and S4. See data availability section for information about availability of the integrative analysis.

We performed our integrative analysis of each cell class individually to maximize the accuracy of cell state mapping across datasets. The seven major cell classes were: excitatory neurons (ExN), inhibitory neurons (InN), astrocytes (Astro), microglia/macrophages (Micro), oligodendrocytes (Oligo), Oligodendrocyte progenitor cells (OPC), and endothelial/pericyte cells (Endo). For ExNs and InNs, we limited our analysis to cortex brain region. However, glial cells were represented from across the brain regions. Table S2 summarizes the datasets that are included in integrative analysis of each of the seven cell classes. As outlined below, we developed a multi-step framework to efficiently handle substantial biological and technical variation that exists among the single cell and nuclei datasets.

#### Selecting highly variable genes

We reasoned that the influence of batch effects on the cell embedding space would be minimized by selection of genes that recurrently show high variability across the human and mouse datasets. To achieve this, we implemented the following method: 1) Select the top 2000 variable genes within each donor of each dataset by the vst method in Seurat^76^. 2) Weight the selected genes in each donor so that the sum of the weights for each dataset add up to one. 3) Calculate an aggregate score for each gene by summing up their weighted scores. This procedure aims to minimize the participation of genes that show between dataset variability (hence likely influenced by batch effects) in the follow up analysis of cell embedding construction and clustering.

#### Principal component analysis

To remove donor-specific batch effects (e.g., due to pre and post-mortem effects, sample preparation, and sequencing settings), we performed scaling (i.e., mean of zero and unit variance) of transcriptome data per gene and per donor and used this scaled data for principal component analysis. Comparison of different integrative solutions indicated the better quality after removal of donor specific effects. For all seven cell class analyses, we used the top 30 principal components, weighted by their variance explained.

#### Batch effect removal

We used Harmony v1.0^77^ to remove batch effects from the PCs with donor id and organism specified as the main source of batch effects. The default theta and lambda parameters were used for all analyses, except for the Endo cells with the theta parameter of four.

#### Assessing the quality integration solutions

To assess the quality of the results, we developed multiple “cluster-free” quality metrics enabling us to perform a systematic and unbiased comparison of the solutions that is independent of the clustering method. These metrics can be grouped into three main categories. First, we required a uniform distribution of the datasets in the UMAP space. In addition to visual inspection, we developed a method that allowed us to do a quantitative evaluation of dataset distributions. Briefly, each of the two UMAP coordinates are split to 100 units, providing 10,000 bins. Within each bin a hypergeometric test is performed to assess whether or not cells from a specific dataset are over-represented. This analysis is performed for each dataset from each integrative solution (the related R functions and visualizations are provided at https://braincelldata.org/resource). Second, we examined whether cells expressing known cortical cell type markers are aligned with each other across datasets and organisms. To systematically test this, we repurposed the commonly used feature plot visualizations to represent donors instead of individual cells, thereby bypassing the effect of sample size variation between donors and datasets (Figure S4). Finally, we assessed if the initial clustering structure of each dataset is preserved in the aligned space. For this analysis, we used the reported clustering structure for each of the datasets individually. We also constructed confusion matrices to compare the cluster annotations between datasets (see the linked website for more details).

#### Cluster quality analysis

On each cell class integrative analysis we performed Leiden clustering using the Seurat package^76^ with clustering resolutions of 0.6 or 0.8. We next used our calculated cell-based quality metrics (nGene, nUMI, MT%, Ribo%, lncRNA%, and %dissociation-related artifact genes) to identify and remove low quality clusters. We also performed marker analysis of each cluster per each dataset using FindAllMarkers() function in Seurat to identify and remove doublet clusters.

#### Uniform annotation of the datasets

We modified a previously developed random walk algorithm^78^ to transfer cell type annotations from the biopsy dataset to each of 35 datasets (27 studies) in the aligned space, thereby uniformly annotating all datasets with cluster labels from the biopsy cohort. We next checked the consistency of cell type proportions among datasets and expression of marker genes across datasets and clusters (Table S3; See the linked website for more details).

### Cell type marker analysis

Marker genes were identified in each dataset by running the ‘FindAllMarkers’ function in Seurat^76^. Significant genes (adjusted p-value < 0.1) with min.diff.pct of 0.1 were considered as markers. The heatmap in Figure 3A based on a select number of microglia markers based on the existing knowledge on microglia states and function. The full organized marker results are provided in Table S3.

To assess an overlap between cell state markers and DE genes in the microglia analysis (Figure 3D), we retained the top 100 upregulated markers (adjusted p-value < 0.01; sorted by p-value) that were markers with logFC >0 in less than a third of the cell states. This additional criterion was added to avoid spurious overlap of markers in non-homeostatic microglia that were driven by the large size of the homeostatic microglia.

### Differential abundance

In our integrative analysis, a major analytic challenge was the wide variation in cell class compositions among the analyzed datasets. As an illustration, both of human PD datasets, one of human AD datasets, and all of mouse AD datasets included only glial cells and not neuronal cells from cortex. To address this, we conducted our meta-analysis of cell type proportional changes within each cell class separately. For our meta-analysis of early AD pathology, we included three out of 6 AD postmortem cohorts from the frontal lobe (more specifically prefrontal cortex or superior frontal gyrus)^3, 4, 6^. We excluded one cohort^79^ due to overlap of individuals with another included cohort^3^, and two that lacked sufficient numbers of early AD stage subjects^5, 7^. We further included two PD postmortem datasets^47, 48^, one ASD dataset^49^ and one MS dataset^50^ as contrast groups in our analyses.

We used a logistic mixed-effect model^80^ to identify differentially abundant cell populations in each dataset separately (Table S4). For all human datasets, we included sex as a fixed effect, and individual as a random effect in the model. We then tested the significance of association between the status with the clusters using a Wald test. For assessing cell type abundance associations with iNPH grading scales, we modified the method to allow for continuous independent variables, while preserving the Wald test for assigning significance.

We performed a meta-analysis of cell type abundance results across four cohorts with designations for earlier and later AD stages: (1) the biopsy cohort (Aꞵ+ as early and Aꞵ+Tau+ as late; all cell classes), (2) Mathys et al. (Braak III-IV as early and Braak V-VI as late; all cell classes except endothelial cells), (3) Leng et al. (Braak III-IV as early and Braak V-VI as late; all cell classes) and (4) Gerrits et al. (CtrlPlus as early and AD as late). The p-values from individual analyses were combined together via the Stouffer’s method, with an additional consideration of the directionality of the change as determined via the odds ratio assessment from the Wald test. All Stouffer’s p-values were subsequently corrected for multiple hypothesis testing via the Benjamini-Hochberg correction.

To determine the relative cellular abundance changes at the cell class level we generated p-values by comparing proportions of the seven cell classes by a Wilcoxon rank-sum test. Meta-analysis p-values (and Z-scores) were generated using Stouffer’s method, taking into account the directionality of the abundance change.

For mouse datasets, we used Fisher’s exact test to examine expansion and loss of different microglia cell types and states (Table S4). The p-values were subsequently corrected for multiple hypothesis testing using the Benjamini-Hochberg procedure. For visualization purposes, z-scores were calculated by transformation of the p-values and signed by the directionality of the log odds ratio.

### Differential gene expression analysis

We employed a pseudocell strategy coupled with mixed linear models and jack-knifing to robustly identify differentially expressed genes. To construct pseudocells, we aggregated the raw UMI count of, on average, every 30 cells per subject and cell type. We constructed one pseudocell for cell types that had between 15 to 45 cells in a donor and excluded cell types that had less than 15 cells. This reduces the impact of dropout and technical variability, while ameliorating low statistical power and high variation in sample size issues attributed to the pseudobulk approaches^81^. We used the Limma Trend^82^ approach with robust moderated t-statistic to identify DE genes within each cell class with sex, cell type, log2(pseudocell MT%) and log2(pseudocell nUMI) as covariates and subject id as a random effect. Cell type annotation was included as a covariate to account for the cell type-specific baseline expression of the genes and therefore to ameliorate the impact of cell type expansion on the DE patterns.

We further performed jack-knife resampling at two levels to identify robust DE genes that are shared among the majority of individuals. First, iterating on each of the 52 individuals in the biopsy cohort, we excluded one subject from the analysis of each cell class and then re-calculated the DE statistic for the remaining 51 individuals, retaining the maximum p-value (i.e., the least significant p-value) achieved for each gene. An adjusted jack-knife p-value was next calculated for genes with adjusted p-value < 0.1 in the main analysis using the Benjamini-Hochberg correction. Second, iterating 50 times, we randomly sampled 50% of cohort subjects (balanced by their pathological status) and re-calculated the logFC patterns. A consistency score was defined for each gene as the fraction of iterations in which the jack-knifed logFCs were consistent with the logFC pattern from the full cohort of 52 individuals (i.e., up- or down-regulated in both). Genes with jack-knifed adjusted p-value < 0.01 and jack-knifed consistency score ≥0.9 were deemed as significant. Comparison of the DE patterns with a pseudobulk approach using LimmaTrend indicated highly consistent (>99%) logFC patterns between the pseudobulk and our pseudocell strategy for the DE genes. We also found majority of identified DE gens by pseudocell approach remain significant (median >72% per cell class; pseudobulk FDR-adjusted p-value < 0.1) in the pseudobulk approach, while the identified DE genes by our pseudocell approach show much less sensitivity to variation in cohort size (data not shown).

To compare the DE genes between Aꞵ+ and Aꞵ+Tau+ individuals (Figure 1F), we performed a paired t-test based on values from below equation:

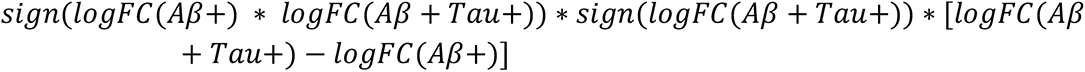

This equation will be positive only if the logFC from both Aꞵ+ and Aꞵ+Tau+ are in the same direction (i.e., gene is up or down regulated in both conditions) and are stronger in Aꞵ+Tau+ and negative otherwise. We excluded from this analysis cell types with less than 20 DE genes in either Aꞵ+ and Aꞵ+Tau+ conditions as they usually were small and their fold change patterns were not reliable. We used a paired t-test to determine if the outputs of this function are randomly distributed around zero or are biased towards positive (i.e., consistent but stronger logFC in Aꞵ+Tau+) or negative (i.e., discordant or stronger in Aꞵ+) values.

### Gene set enrichment analysis

DE genes were filtered to protein-coding based on the gene biotype information from the ‘EnsDb.Hsapiens.v86’ package in R Bioconductor. Genes were next ordered based on their t-statistic from LimmaTrend mixed linear models. Curated GO Biological Process, KEGG and Reactome gene sets were retrieved from the EnrichR portal^83^. To identify enriched pathways, we ran the fGSEA package^84^ v1.16.0 with default setting while limiting the geneset sizes between 15 and 250 genes. For each cell class and cell type analysis, protein-coding genes expressed in more than one percent of cells in the corresponding group were used as background.

### Correlating cell type proportional changes to transcriptome responses across cell types

To identify correlation between cell type abundances and transcriptional phenotypes, we used a logistic mixed-effect model^80^. Specifically, we constructed a meta-gene (referred to as DE signature in the main text) from the top 300 upregulated protein-coding genes (sorted by their jack-knifed p-value) in each individual cell by aggregating their corresponding UMI counts, as proposed before^85^. The meta-gene was next normalized over the total nUMI count of the cells and standardized to have a mean of zero and variance of one across cells from all subjects in our cohort. We then binarized cells as active or inactive for a meta gene based on a standardized score threshold of 2. In our analyses we required presence of at least two subjects with more than 3% transcriptionally active cells in each cell type and status category. For cell types that do not meet this criterion we set the association p-value to one to indicate transcriptional changes in the corresponding cell type are not associated with the interested cell type fractional variation. Finally, we fit a logistic mixed-effect model on the binarized scores to examine their association with the normalized cell type proportional changes with sex as a covariate and subject as a random effect. The cell type proportions were normalized by applying an empirical cumulative estimation using the ecdf() function in R. We used cell-class level DE genes to construct meta-genes since: 1) cell class level DE genes were highly conserved within cell types (Figure S6D); 2) DE genes were not driven by the variation in the cell counts of the cell types. As summarized below, we performed robustness analysis and alternative meta-gene construction schemes to further confirm the observed associations.

First, to assess the robustness of the results, we tested the sensitivity of the results to the presence of technical variation in cell gene counts by, iterating 30 times, adding Poisson noise to the transcriptome data of each individual cell before calculation of the meta-gene expressions (Figure S7B). In addition, iterating 30 times we randomly down sampled the cell types to examine the association of the cell type sizes on the results (Figure S7C).

Second, as an alternative analysis method to support our findings (Figure S7D), we constructed meta-genes through principal component analysis of the normalized and scaled expression of the top 300 upregulated protein-coding genes (sorted based on jack-knifed p-values). Similar to the WGCNA approach^86^, the first principal component was chosen as the meta-gene. The meta-gene scores were then binarized as above and the association with cell type proportional changes were examined using a logistic mixed-effects model similar to above.

### Heritability enrichment of differentially expressed genes with MAGMA

We used MAGMA^42^ to determine the degree of enrichment of common variant risk in the list of differentially expressed genes across cell types. We first downloaded the summary statistics from a recent common variant meta-analysis of AD and related dementias^35^, PD^87^, and ASD^88^. Using the online FUMA tool^89^, we generated Z-scores for each gene, corresponding to the approximate degree of association between the gene and AD (SNP2GENE function). To determine gene sets for each cell class, we took the top 200 differentially expressed genes between biopsy samples with AD pathology versus those samples without, ordered by t-statistic at a significance value of p < 0.2 (to ensure enough genes were being captured per gene set). Significance values for the gene set of interest were calculated via MAGMA, wherein a regression is fit to determine whether those genes with membership for that set have a significant enrichment for heritable risk of the trait of interest.

### Generation of a doxycycline-inducible *SOX10* H1 stem cell line

We adapted a recently-published protocol to produce mature oligodendrocytes from the H1 embryonic stem cell precursors cell line^90^. First, we isolated and incorporated a *SOX10* transcription factor (Addgene #115242) into the backbone of a doxycycline-inducible cassette (Addgene #105840) to generate pBR01.

H1 ESCs were plated in Matrigel (cat. #47743-716)-coated (30-minute incubation at 37°C prior to cell plating) plates in mTesR1 (cat. #85857) with supplements (ESC media, StemCell Technologies cat. #85857) and RevitaCell (cat. #A2644501). After plating, we performed daily media changes with ESC media without RevitaCell until plates were approximately 80% confluent with compact colonies. For routine passaging, ESCs were washed 1X with PBS (cat. #10010049) and incubated in Versene (cat. #BE17-711E) for 5 mins at room temperature, after which Versene was gently aspirated from the plate and replaced with ESC media. ESCs were gently dissociated into a cell suspension using a manual cell scraper and transferred as small colonies to a fresh Matrigel-coated plate at a 1:20 dilution. In order to generate a doxycycline-inducible *SOX10* cell line, we performed TALEN-based integration as has been previously described^91^. Briefly, we electroporated (1050V, 50 ms pulse, two pulses total) 1 million ESCs with 4 μg pBR01, 2 μg TALEN-L (Addgene #59025), 2 μg TALEN-R (Addgene #59026), and 0.4 μg Bcl-XL (Addgene #8790) plasmids using a Neon Transfection System Pipette Station (Thermo Fisher). After 96 hours, cells were incubated with 2 μg/mL puromycin (cat. #A1113803) for 72 hours. After puromycin selection, polyclonal ESCs were expanded and stored in liquid nitrogen at 10^6^ cells/mL in 10% DMSO + mTesR1.

### Oligodendrocyte differentiation of ESCs

To differentiate the resulting cell line into mature oligodendrocyte lineage cells, we adapted the Garcia-Leon protocol with minor modifications^90^. Briefly, we used the H1 embryonic stem cell line with the integrated *SOX10* cassette to generate neural progenitor cells which were subsequently differentiated into mature oligodendrocytes. Vials containing 1 million ESC precursors were thawed and plated into one well of a Matrigel-coated 6-well plate, supplementing the cells with RevitaCell to increase vitality. These cells were then allowed to grow to confluence with supplementation of 1mL E8 media (cat. #A2858501). Confluent cells were subsequently split with the following procedure. First, cells were washed with 1mL of Dulbecco’s PBS per well. Then, cells were subsequently treated with 1mL of ReleSR (cat. #05872) incubated at 37°C for five minutes. Cells were then spun down and split at a ratio of 1:14 into Matrigel-coated 6-well plates and grown in 1mL of E8 media (cat. #A2858501) supplemented with 1x RevitaCell solution. The cells were transitioned to mTeSR1 media by replacing with a 1:1 E8 to mTeSR1 solution on the first day, 75% mTeSR1 with 25% E8 on the second day, and a full 1mL of 100% mTeSR1 on the thirdday. Cells were then allowed to grow to confluence before being split again with 1mL ReleSR as above. The cells were replated onto 6-well matrigel-coated plates, supplemented with RevitaCell. An N2B27 media was made by mixing non-essential amino acid MEM (cat. #11140-050), 2-mercaptoethanol (cat. #21985023), N2 (cat. #17502048) and B27 (cat. #12587010) to 1× concentration plus insulin (cat. #I9278) at 25 µg/ml final concentration to Dulbecco’s modified essential media. The cells were then grown in 2 mL of the pre-made N2B27 medium supplemented with 0.1 μM retinoic acid (RA, cat. #R2625), 10 μM SB431542 (cat. #04-0010-10) and 1 μM LDN193189 (cat. #04-0074) for five days and an additional two days with 10 μM of smoothened agonist (SAG, cat. #566660).

After cells achieved confluence, they were passaged using a pre-warmed 1mL aliquot of Accutase (cat. #A1110501) for 1-2 minutes. The cells were seeded onto 6-well plates coated with poly-l-ornithine (cat. #P3655) and laminin (cat. #L2020-1MG)-coated plates at a density of 10,000 cells per square centimeter. The cells were fed a differentiation medium supplemented with 2 μg/mL of doxycycline (cat. #D9891-10G) to induce expression of *SOX10* and allowed to grow for 10 days, at which time they are mature.

### Neuronal differentiation of ESCs

Neuronal differentiation of ESCs into cortical glutamatergic neurons was carried out as previously described^92^. In brief, the differentiation was carried out by adding doxycycline hyclate (2 μg/mL) to N2 supplemented media (Thermo Fisher, 17502048) with patterning factors SB431542 (Tocris, 1614, 10 μM), XAV939 (Stemgent, 04-00046, 2 μM) and LDN-193189 (Stemgent, 04-0074, 100 nM), as described previously^92–94^. Puromycin selection was used (5μg/μL), from days 2 to 6 to remove non-transduced cells. At 4 days post induction, neuronal cells were resuspended into Neurobasal media (Gibco, 21103049) that was supplemented with B27 (Gibco, 17504044, 50X), doxycycline (2 μg/mL), brain-derived neurotrophic factor (BDNF), ciliary neurotrophic factor (CTNF), and glial cell-derived neurotrophic factor (GDNF) (R&D Systems 248-BD/CF, 257-33 NT/CF, and 212-GD/CF at 10 ng/mL each). From this point onwards the neurons were either co-cultured with murine glial cells that were derived from early postnatal (P1-P3) mouse brains as described previously^95^ or were left to grow as monocultures (mouse strain https://www.jax.org/strain/100012; animal ethical committee approval by Harvard University: Animal Experimentation Protocol (AEP) # 93-15).

### Immunohistochemistry and imaging of ESCs

We performed immunohistochemistry on ESC-derived oligodendrocytes at 1, 5, and 10 days after doxycycline-based *SOX10* induction. Briefly, cells were grown on a 6-well plate and fixed using 2% PFA and then permeabilized using Triton-X (cat.# T9284-1L), followed by multiple washes with 1x Dulbecco’s PBS at each step. We used the following primary antibodies for our immunohistochemistry experiments: anti-*O4* (cat. #MAB1326), anti-*MBP* (cat. #AB9348), anti-*NeuN* (cat. #MAB377), and anti-*PAX6* (cat. #AB78545). The primary antibodies were diluted in a solution of 10% bovine serum albumin (BSA) in phosphate-buffered saline (PBS) supplemented with 1% Triton-X then added to the cells and allowed to incubate overnight at 4°C. Cells were then washed three times in PBS. Finally, the secondary antibodies were diluted in a solution of 10% BSA in PBS supplemented with 1% Triton-X then added to the cells and allowed to incubate for 1-2 hours at room temperature. After the secondary incubation, one to two drops of ProLong Glass AntiFade Mountant with NucBlue (cat.# P36981) was added into the wells and coverslips were added on top of each cell culture into wells for downstream imaging.

Imaging of immunohistochemical stains was performed on a Keyence BZ-800XE microscope with a Nikon Apo 20x objective. All images were acquired using the same light emission settings and all channels were set to the same LUTs before quantification. For quantification, we used CellProfiler’s IdentifyPrimaryObjects and MeasureObjectIntensity function to segment cells based on their DAPI signal. Subsequently, the average fluorescence value (mean intensity value) was normalized per cell to the average fluorescence intensity of the DAPI signal. To determine the significance of an intensity difference, a linear mixed-effect model was used to calculate the significance of a change in normalized intensity value across days of differentiation, treating each slice image as a random effect. Significance values were determined via a likelihood ratio test against the null model not containing the day of differentiation.

### Generation of single-cell suspension from ESC-derived H1 iOligodendrocytes

To generate single-cell experiments, briefly we used oligodendrocytes at terminal differentiation (past day 8 post-doxycycline induction of SOX10). We isolated cells using the passaging protocol as mentioned above and measured cell concentrations in our isolate using a hemocytometer.

### Read processing and clustering of iOligodendrocyte and iExcitatory Neuron scRNA-seq experiments

Sequencing reads from iOligo experiments were demultiplexed and aligned to the hg38 reference using CellRanger with default setting using the command CellRanger mkfastq, followed by count generation using the command CellRanger count. Sequencing reads from iExN experiments were demultiplexed and aligned to the hg38 reference using DropSeqTools with default setting.

To analyze single-cell RNA-sequencing data from ESC-derived oligodendrocytes and neurons we first determined highly variable genes using LIGER. We further used non-negative matrix factorization (with k, number of factors, set to 20) to determine a low-dimension embedding followed by graph-based clustering using SLM. Marker genes were identified by a Wilcoxon rank-sum and cells were annotated based on known markers of mature cell types as identified from our biopsy dataset.

### ELISA-based amyloid beta quantification

To quantitate amyloid beta peptide levels from cell culture, we used the MesoScale Discovery V-Plex Plus Aꞵ Peptide Panel 1 (6E10) ELISA kit (cat. #K15200G). Briefly, we extracted 1.5 mL of conditioned media per well replicate from isolates of ESC-derived oligodendrocytes, neurons, and microglia. Isolates were stored at -80°C till the ELISA assay was run at which time they were brought up to 4°C before being spun down at 10,000rpm for 15 minutes. The MSD ELISA was run according to the manufacturer’s guidelines. Absolute Aꞵ peptide abundances were quantified using the MSD Discovery Workbench Analysis Software.

---

**Figure S1.**
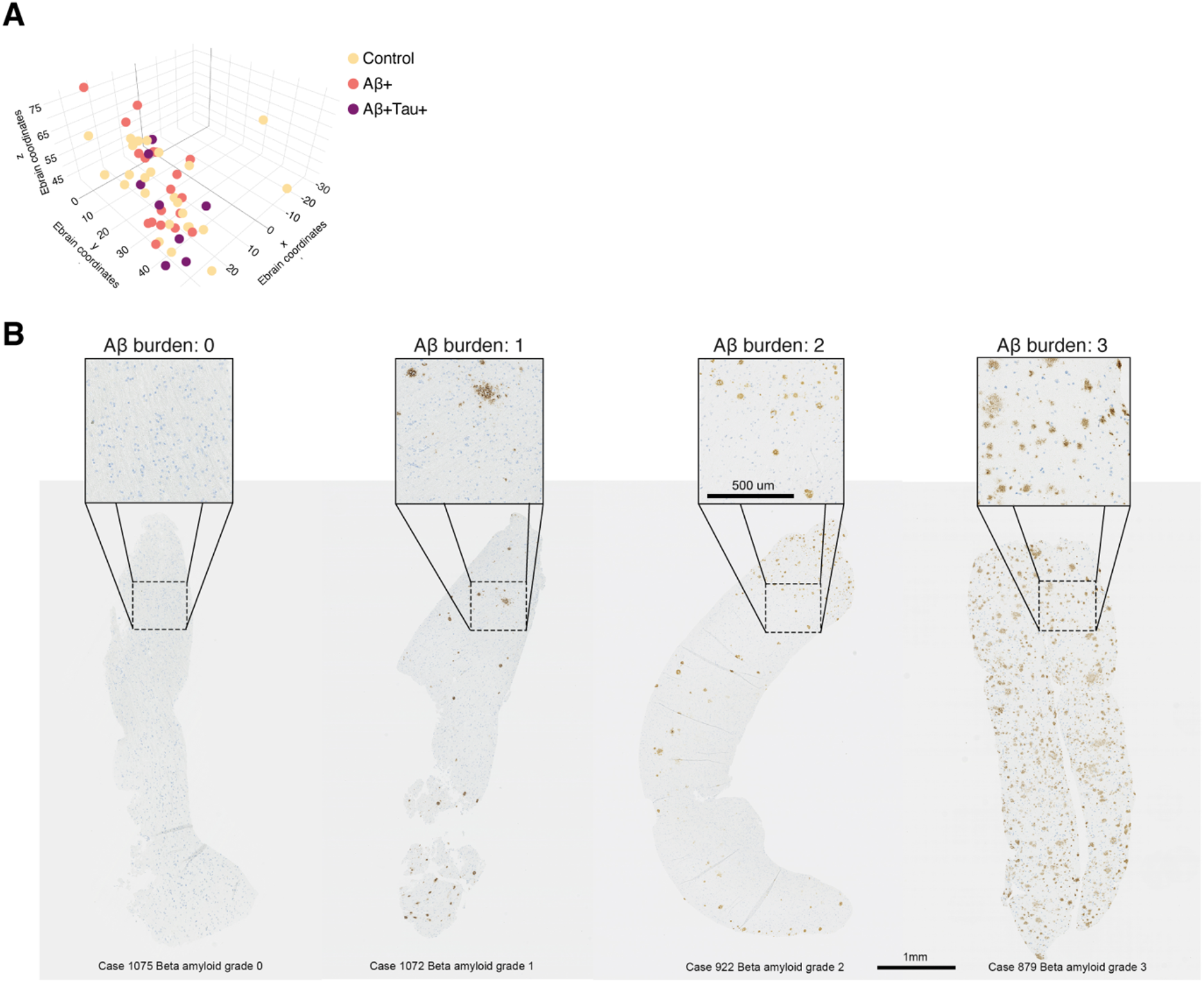
Stereological positioning and neuropathological scoring of biopsy cohort samples. ***A)*** *Relative three-dimensional coordinates of biopsy positions based on the mapping of post-surgical CT or MRI images from 52 subjects. Samples are colored by AD pathologic status. **B)** Representative images of biopsies with different Aꞵ burden*.

**Figure S2.**
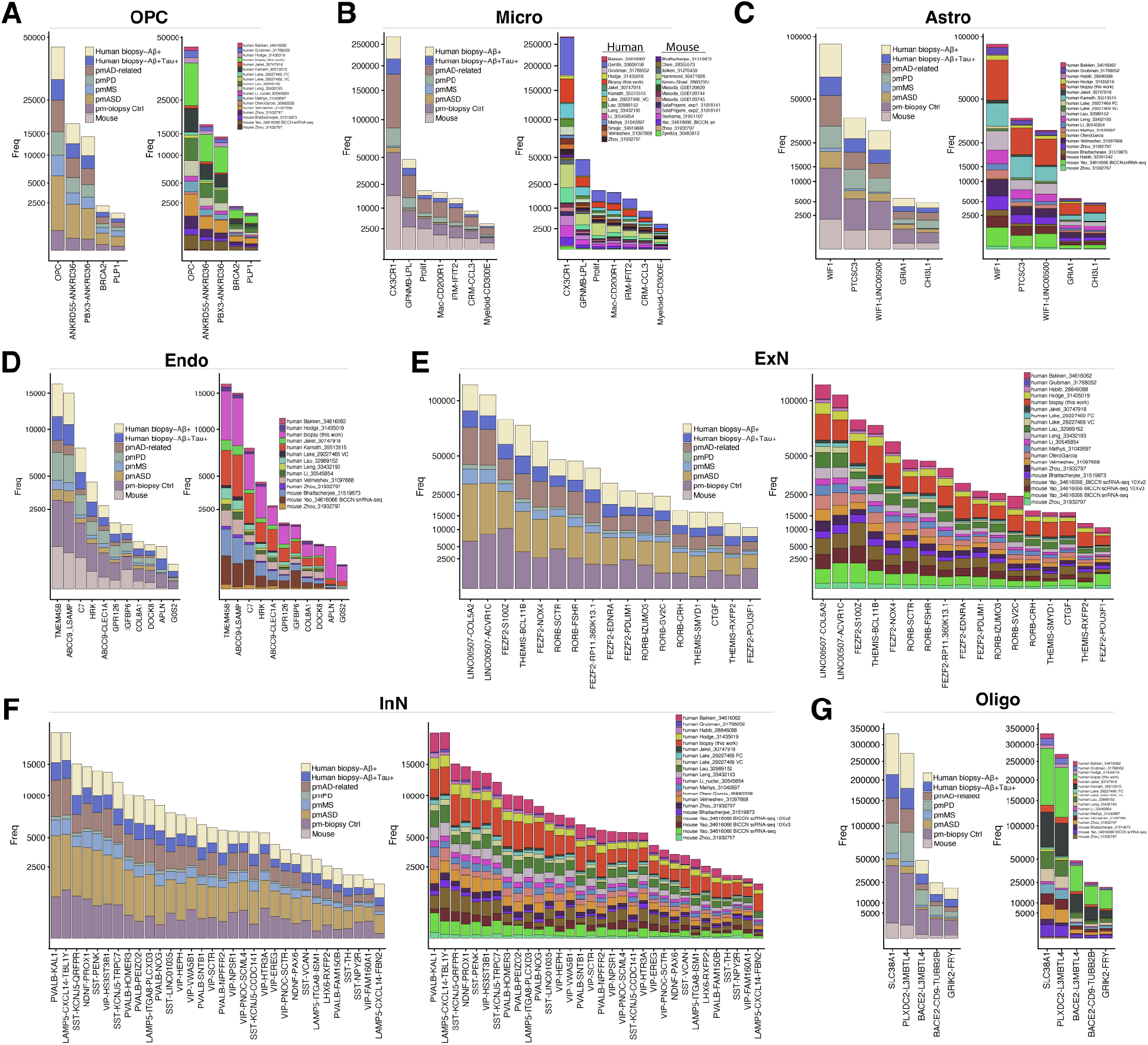
Individual cell types in the integrative analysis are well represented across biological conditions and datasets. ***A-G)*** *Number of cells in each cell type stratified by biological condition (left), and dataset (right). pm, postmortem; PD, Parkinson’s disease; MS, multiple sclerosis; ASD, autism spectrum disorder*.

**Figure S3.**
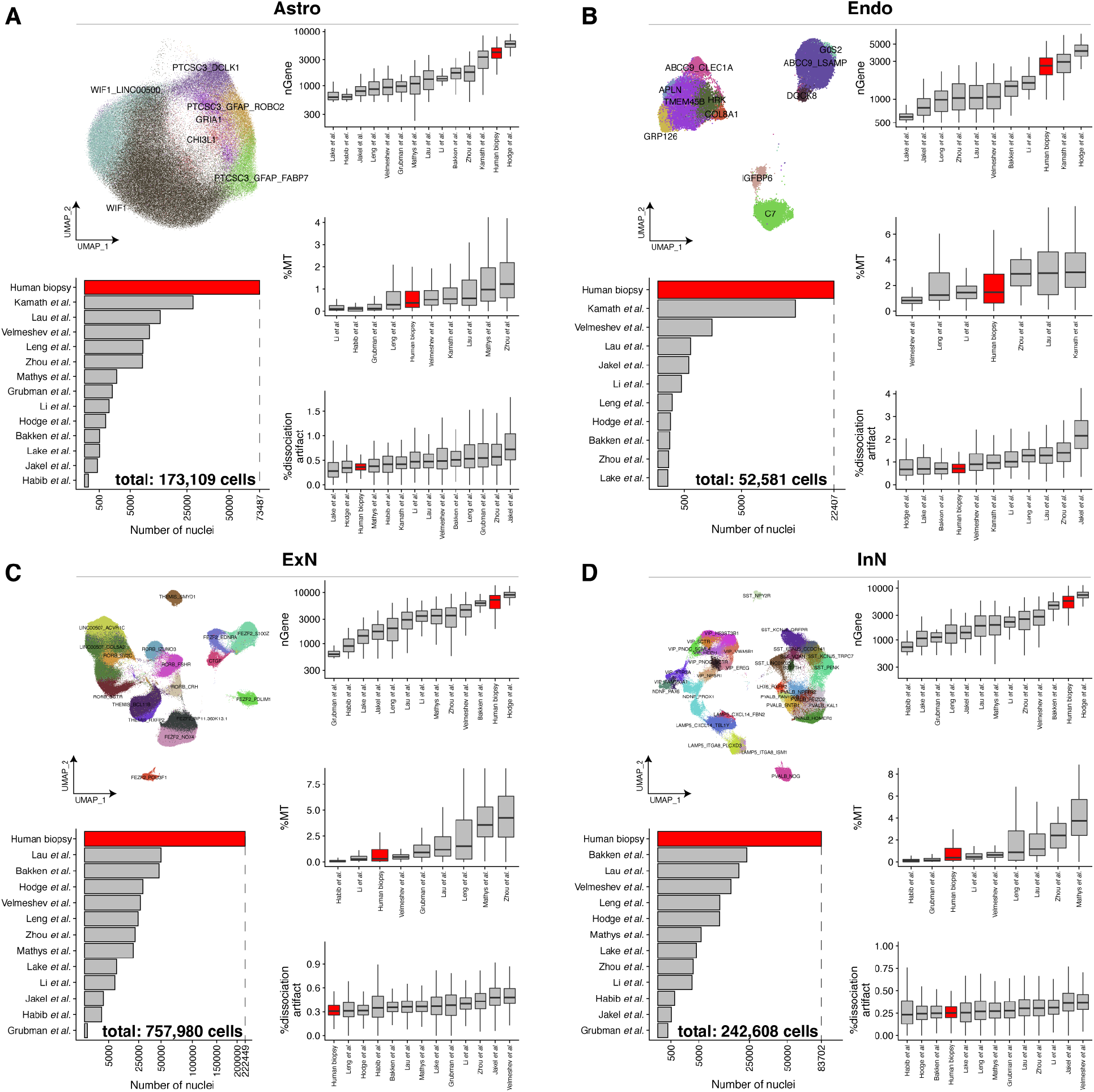
Quality metrics of human datasets included in the integrative analysis of Astro, Endo, ExN, and InN cell classes. ***A-D)*** *QC metrics of human datasets parsed by major cell class: A) Astrocytes, B) Endothelial cells, C) Excitatory neurons, D) Inhibitory neurons. Detailed marker analysis results are provided in Table S3. More QC metrics are available through the linked portal (see data availability section). %MT: percent expression of genes per cell that map to mitochondrial genes, normalized by nUMI. %dissociation artifact: percent expression of dissociation-related artifactual genes*^20^ *in each nuclei and normalized by nUMI. Astrocyte PTCSC3 cell type is more deeply clustered into three states (PTCSC3-DCLK1,PTCSC3-GFAP-ROBO2, and PTCSC3-GFAP-FABP7). Center line, median; box limits, upper and lower quartiles; whiskers, 1.5x interquartile range*.

**Figure S4.**
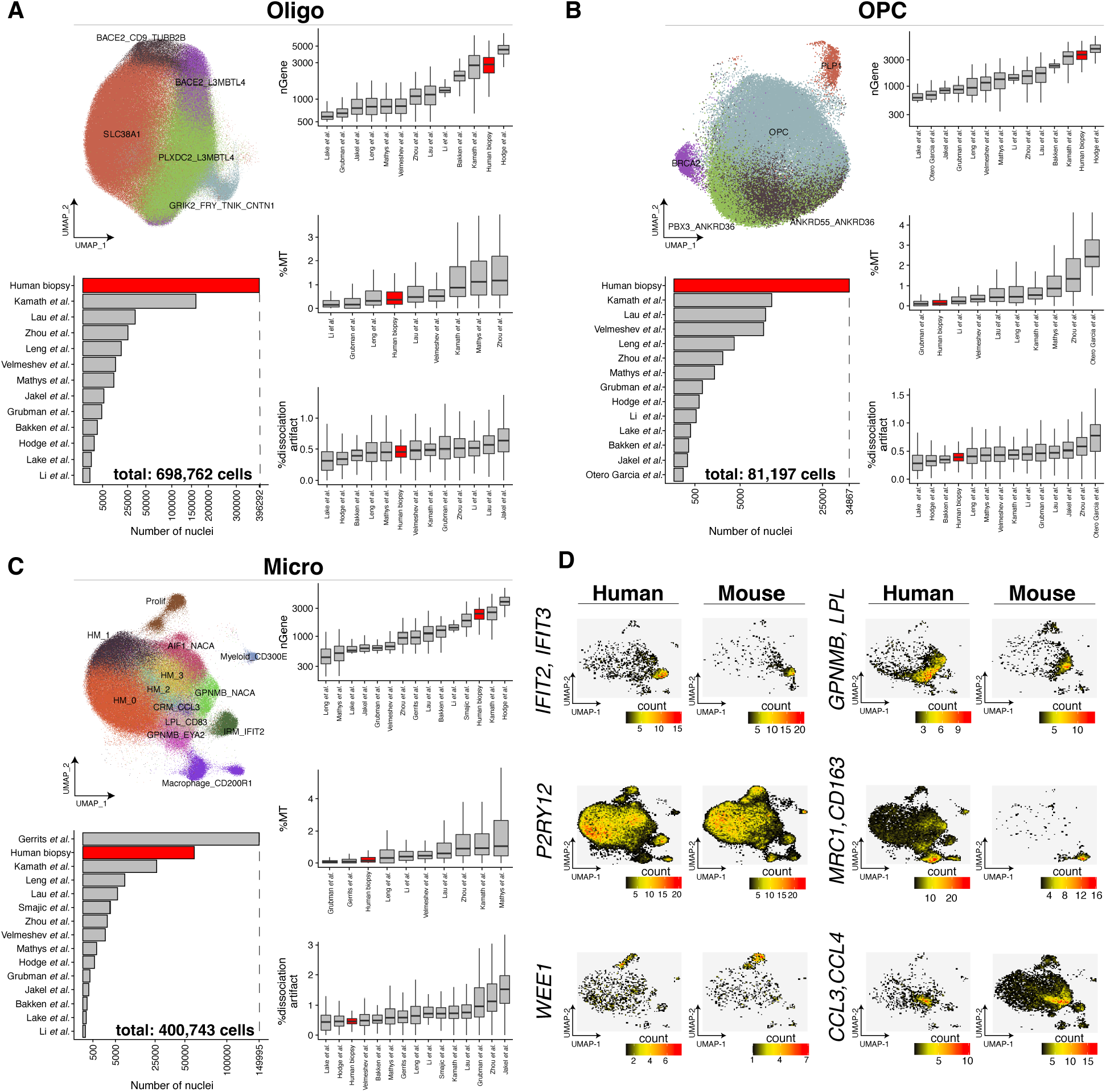
Quality metrics of human datasets included in the integrative analysis of Oligo, OPC, and Micro cell classes. ***A-D)*** *QC metrics of human datasets parsed by the major cell class: **A)** Oligodendrocytes, **B)** Oligodendrocyte progenitor cells, and **C)** Microglia. **D)** Comparisons of some key microglia states between human and mouse datasets. Color indicates the number of human or mouse donors that support the expression of the gene in a given UMAP coordinate across datasets. Detailed marker analysis results are provided in **Table S3**. More QC metrics are available through the linked portal (see data availability section). %MT: percent expression of mitochondrial genes in each cell and normalized by total nUMI. %dissociation artifact: percent expression of dissociation-related artifactual genes in each cell and normalized by total nUMI. Microglia cell types CX3CR1 and GPNMB-LPL are more deeply clustered into five (HM-0 to HM-4) and three (LPL-CD83 ,GPNMB-EYA2, GPNMB-NACA) states, respectively. Center line, median; box limits, upper and lower quartiles; whiskers, 1.5x interquartile range*.

**Figure S5.**
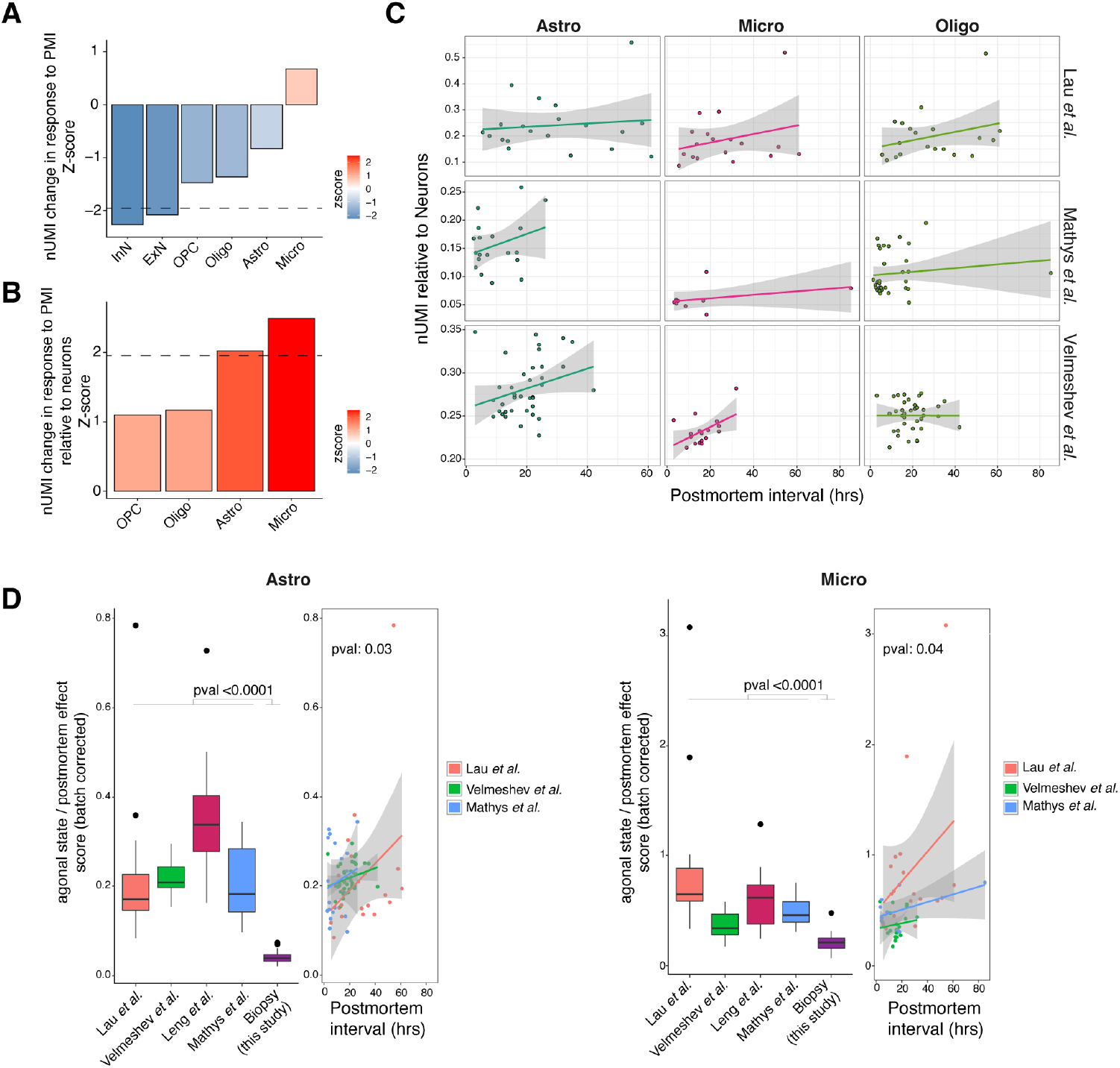
Peri- and post-mortem effects on gene expression patterns. ***A)*** *Impact of postmortem interval on the number of expressed transcripts (nUMI) per cell class. To estimate significance within each postmortem dataset, a regression line was fit to estimate the significance of association between PMI and the mean nUMI in each subject. For this analysis, we considered three postmortem datasets of Lau et al., Mathys et al., and Velmeshev et al. that their postmortem interval information were available and had sufficient range for a regression analysis. The p-values from each dataset were next combined using Stouffer’s method. To calculate the nUMIs per cell, we excluded the top 50 expressed genes in each dataset to better capture the impact of the postmortem intervals on the expression of the lower expressed genes. The dashed line represents the p-value cutoff threshold of 0.05. **B)** Impact of PMI on glial-to-neuronal gene expression ratio. Within each of three postmortem datasets, Similar to panel A, a regression line was fit to examine the impact of the PMI on the mean ratio of glial-to-neuronal genes in each subject. The p-values were next combined using Stouffer’s method. In each subject, the mean neuronal expression was calculated as the mean nUMI of excitatory and inhibitory neurons, excluding the top50 expressed genes. The dashed line represents the p-value cutoff threshold of 0.05. **C)** Glial-to-neuronal gene expression ratio as a function of PMI in each of the three postmortem datasets. **D)** Comparison of glial-to-neuronal gene expression levels between the biopsy and four postmortem datasets. Glial expression in each dataset was normalized to reduce the effect of the ambient RNA as measured by the expression level of the top 250 most specific markers of excitatory and inhibitory neurons. The neuronal specific gene markers were identified based on the pct.1 - pct.2 difference in our biopsy dataset. As shown, this normalization resulted in overlay of agonal state/postmortem scores for samples with a similar PMI across datasets*.

**Figure S6.**
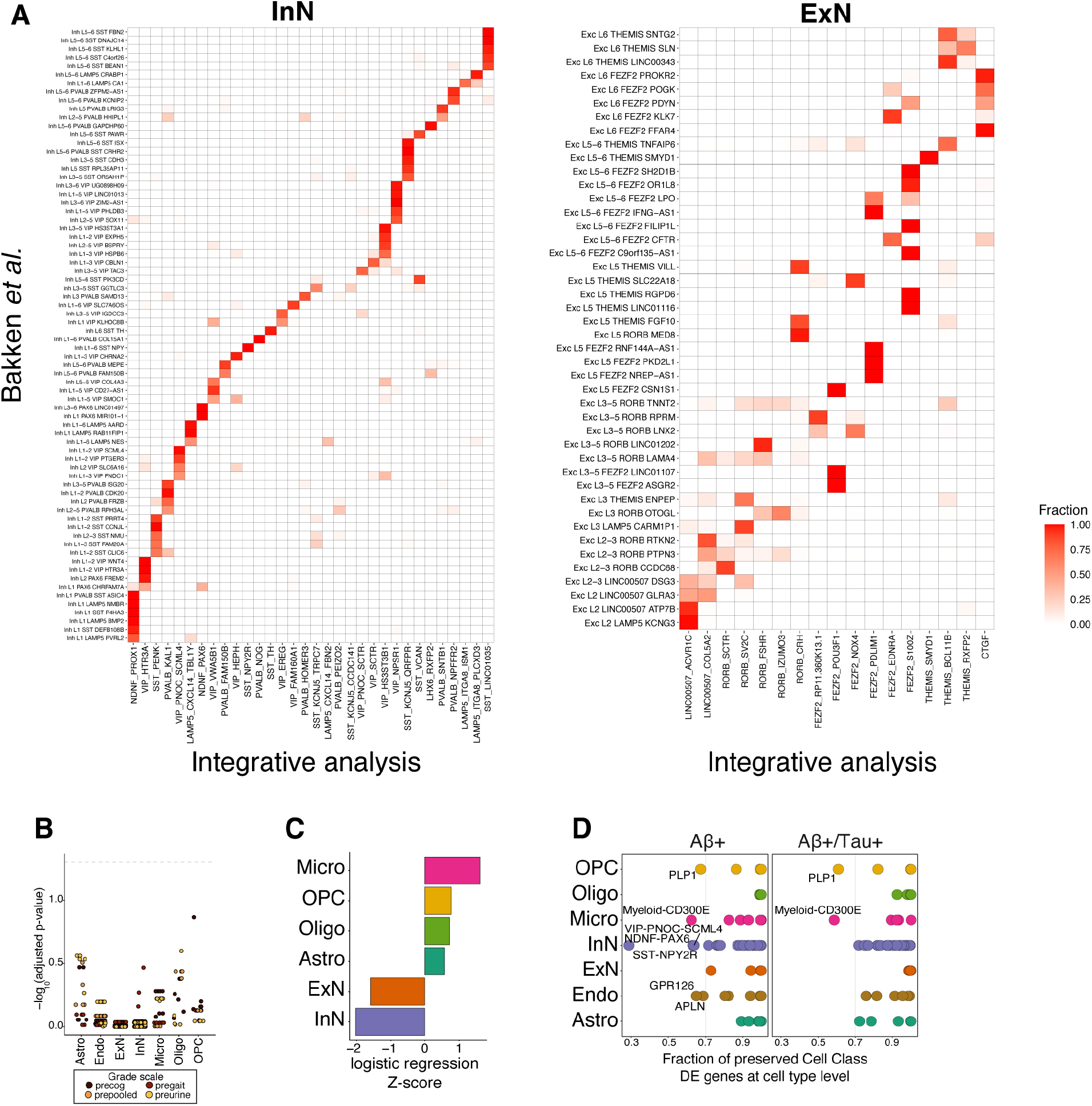
Correspondence of neuronal cell types from integrative analysis with a reference dataset containing cortical layer annotations. ***A)*** *The plots represent the correspondence of neuronal cell type annotations from our integrative analysis with cell cluster annotations from a human cortex dataset*^21^ *that included cortical layer information. The sum of fractions in each row adds up to one. B) Dot plot of -log_10_-transformed FDR-adjusted p-values from logistic mixed-effect model testing association of cell type abundance with iNPH GS (Methods) subscore severity measured prior to shunt placement (precog = cognitive subscale, pregait = gait subscale, preurine = urine subscale, prepooled = combined iNPH GS score). C) Z-score from meta-analysis of cell class level differential abundance comparing late-stage samples (Braak V-VI and Aꞵ+Tau+) versus pathology-free samples (Methods). D) Cell class level DE genes are preserved within cell types. Cell types are labeled in which less than 70% of cell class level DE genes were preserved*.

**Figure S7.**
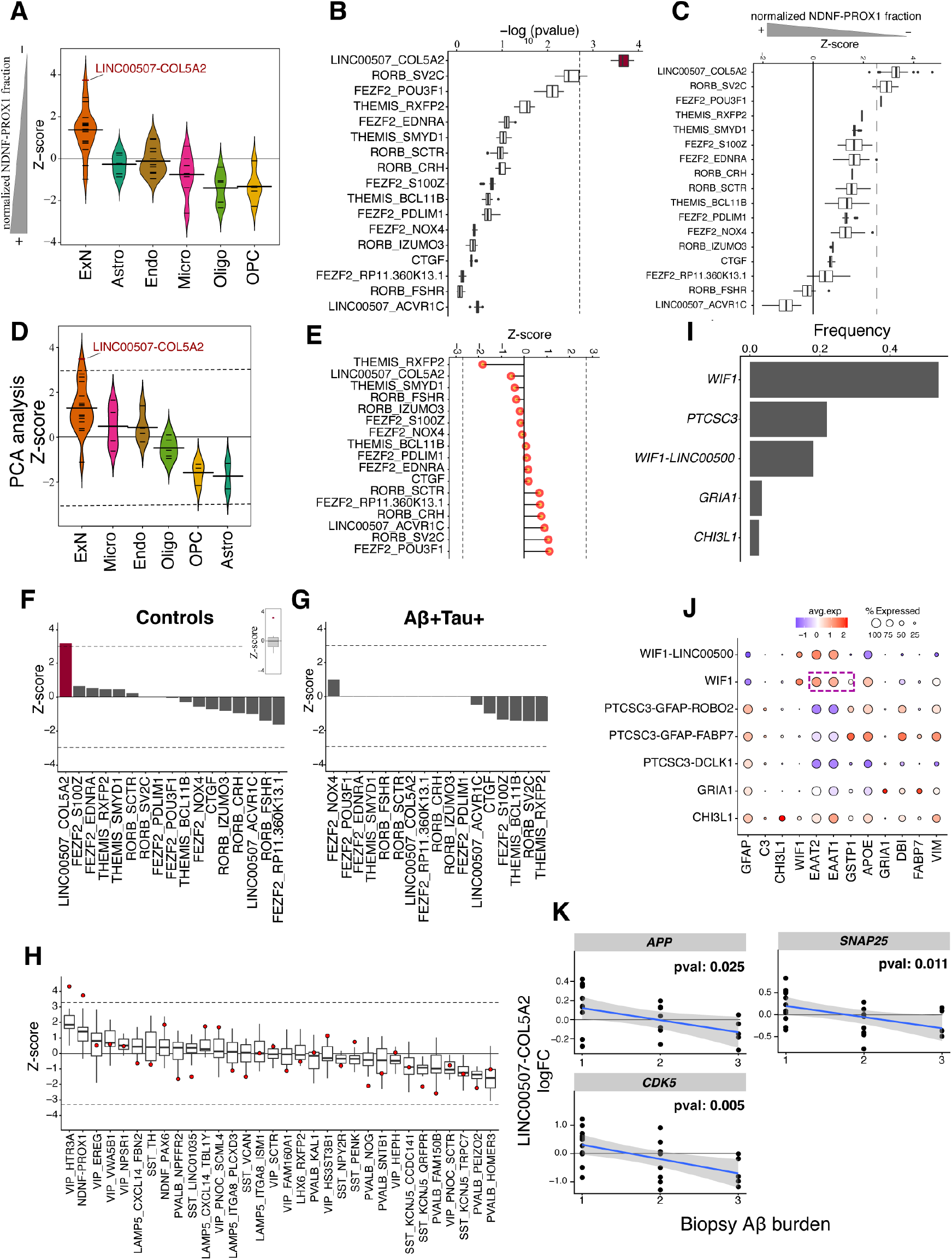
Loss of NDNF-PROX1 inhibitory neurons is associated with an upregulation of excitatory neuron DE signature. ***A)*** *Distribution of Z-scores per cell class for the analysis shown in* Figure 3A*. Each small line indicates one cell type and the tick lines represent the mean. **B)** Logistic mixed-effect model regression (Methods) of NDNF-PROX1 proportion versus ExN cell type transcriptional signature in Aꞵ+ subjects with added poisson noise. Poisson noise counts were added to the UMI counts of each gene in each cell prior to computing the regression. Boxplots show the distribution of -log_10_ transformed p-values over 30 noise iterations. Center line, median; box limits, upper and lower quartiles; whiskers, 1.5x interquartile range; points, outliers. **C)** Logistic mixed-effect model regression of NDNF-PROX1 proportion versus ExN cell type transcriptional signature in Aꞵ+ subjects with downsampling of cells. Iterating 30 times, we randomly downsampled each ExN type to 7000 cells, unless the cell type size was less than this number. Boxplots show the Z-score distributions over the 30 downsampling iterations. The dashed line indicates the FDR threshold of 0.05. Center line, median; box limits, upper and lower quartiles; whiskers, 1.5x interquartile range; points, outliers. **D)** Association between NDNF-PROX1 loss and LINC00507-COL5A2 cell type assessed using an alternative strategy of constructing a meta gene of the ExN DE signature from the first principal component (Methods). Each small line indicates one cell type and the tick lines represent the mean. **E)** Logistic mixed-effect model regression of NDNF-PROX1 proportion versus ExN cell type transcriptional signature in Aꞵ+ subjects after randomizing assignment of cells to excitatory cell types. Dashed line represents FDR-threshold of 0.05. **F-G)** Logistic mixed-effect model regression (Methods) of NDNF-PROX1 proportion versus ExN cell type transcriptional signature in control (**F)** and Aꞵ+Tau+ (**G**) subjects. The inset boxplot in **F** shows the overall distribution of Z-scores among the 17 excitatory neuron types. Dashed line represents FDR-threshold of 0.05. **H)** Boxplots representing the association (as measured by Z-score from logistic mixed-effect model regression) between each inhibitory neuron cell type (x-axis) with the ExN DE signature across 17 ExN types (boxplots). The red dot represents the Z-score of the LINC00507-COL5A2 type. Analysis is based on Aꞵ+ individuals only. In boxplots, center line, median; box limits, upper and lower quartiles; whiskers, 1.5x interquartile range. **I)** Barplots representing the relative frequencies of the five astrocyte cell types. **J)** Dot plots showing expression of astrocyte marker genes in each astrocytic cell type. **K)** logFC expression of APP, CDK5, and SNAP25 genes in LINC00507-COL5A2 excitatory neurons across increasing Aꞵ burden scores. Regression line is illustrated in blue with associated standard error*.

**Figure S8.**
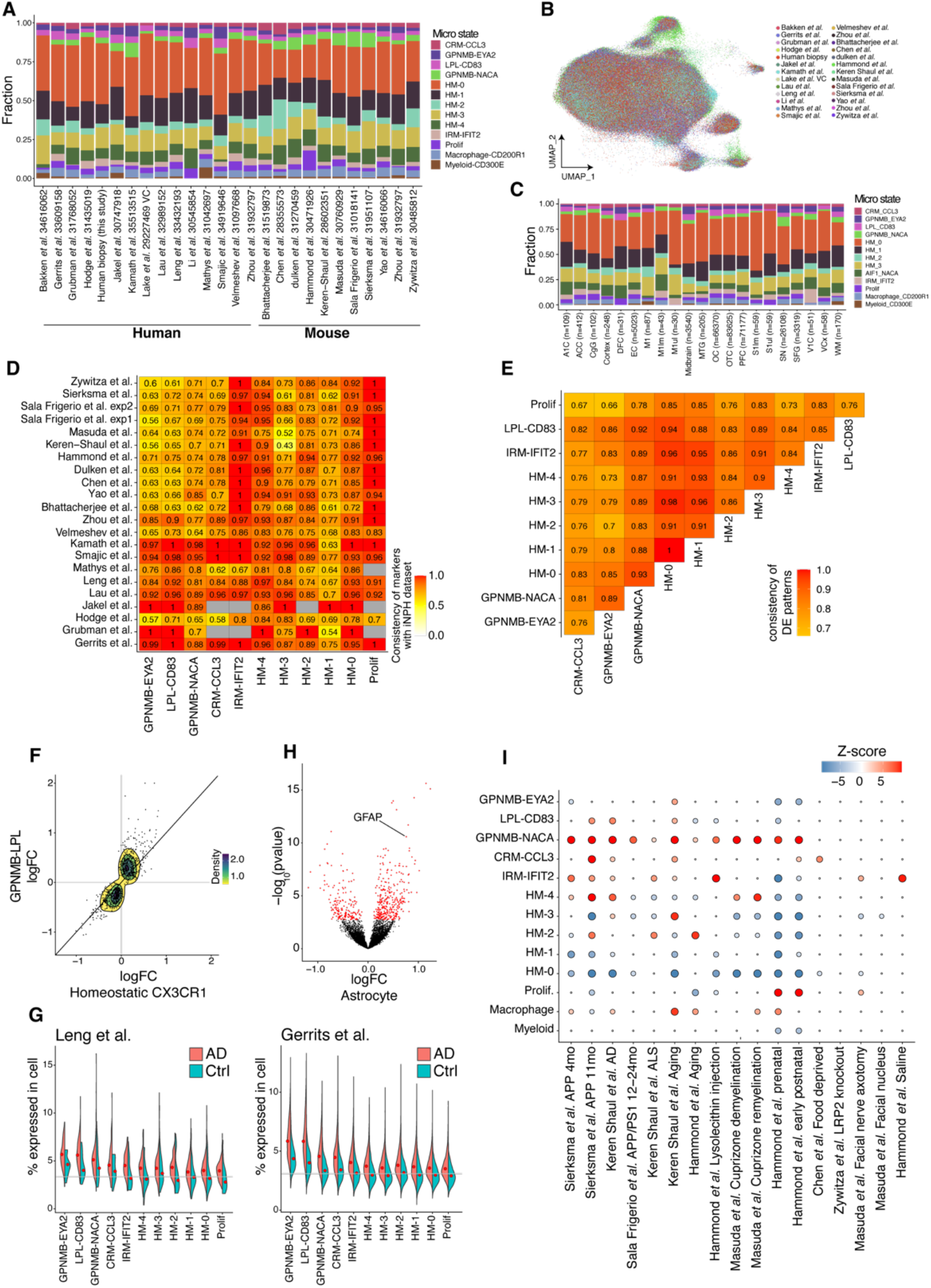
Microglia responses to the accumulation of Aꞵ and tau in cortical tissue. ***A)*** *Microglia cell state compositions across human and mouse datasets. **B)** UMAP representation of microglia integrative analysis where each cell is colored by its dataset of origin. **C)** Proportion of microglial states stratified by brain region. **D)** Marker expression consistency of previous human and mouse single-cell datasets with the biopsy dataset. Consistency score is defined as the fraction of markers from human biopsy dataset for each microglia state that show a conserved up- or down-regulation pattern in each dataset. **E)** The fraction of DE genes from the biopsy dataset between each pair of microglia states that have a conserved logFC pattern (e.g., up or down in both cell states). DE genes were calculated by comparing Aꞵ+/Aꞵ+Tau+ samples with controls. **F)** Comparison of DE genes between GPNMB-LPL and CX3CR1 microglia types in the biopsy dataset. The analysis is based on the union of top 300 DE genes in each cell type to reduce the impact of cell type size variation (i.e., statistical power). **G)** Microglia DE genes from the biopsy dataset are upregulated in two AD postmortem studies and are enriched for the markers of GPNMB-LPL and LPL-CD83 microglia states. Meta-gene expression of the up-regulated microglial DE genes from our dataset in two published postmortem studies*^4, 6^*. The meta-gene was constructed by summing their UMI counts in each cell and normalizing by the nUMI. The gray lines illustrate the median expression of the meta-gene across microglia states. **H)** Differential expression analysis of astrocyte genes in response to Microglia GPNMB-EYA2 and LPL-CD83 expansion. The DE genes (FDR-adjusted p-value < 0.05) are represented in red. Fractions of microglia GPNMB-EYA2 and LPL-CD83 cells in individuals are normalized using an empirical normal cumulative estimation function (ecdf function in R) to have a range between zero and one. **I)** Enrichment of mouse microglia states in response to various conditions (Fisher’s exact test; FDR-adjusted p-value < 0.1)*.

**Figure S9.**
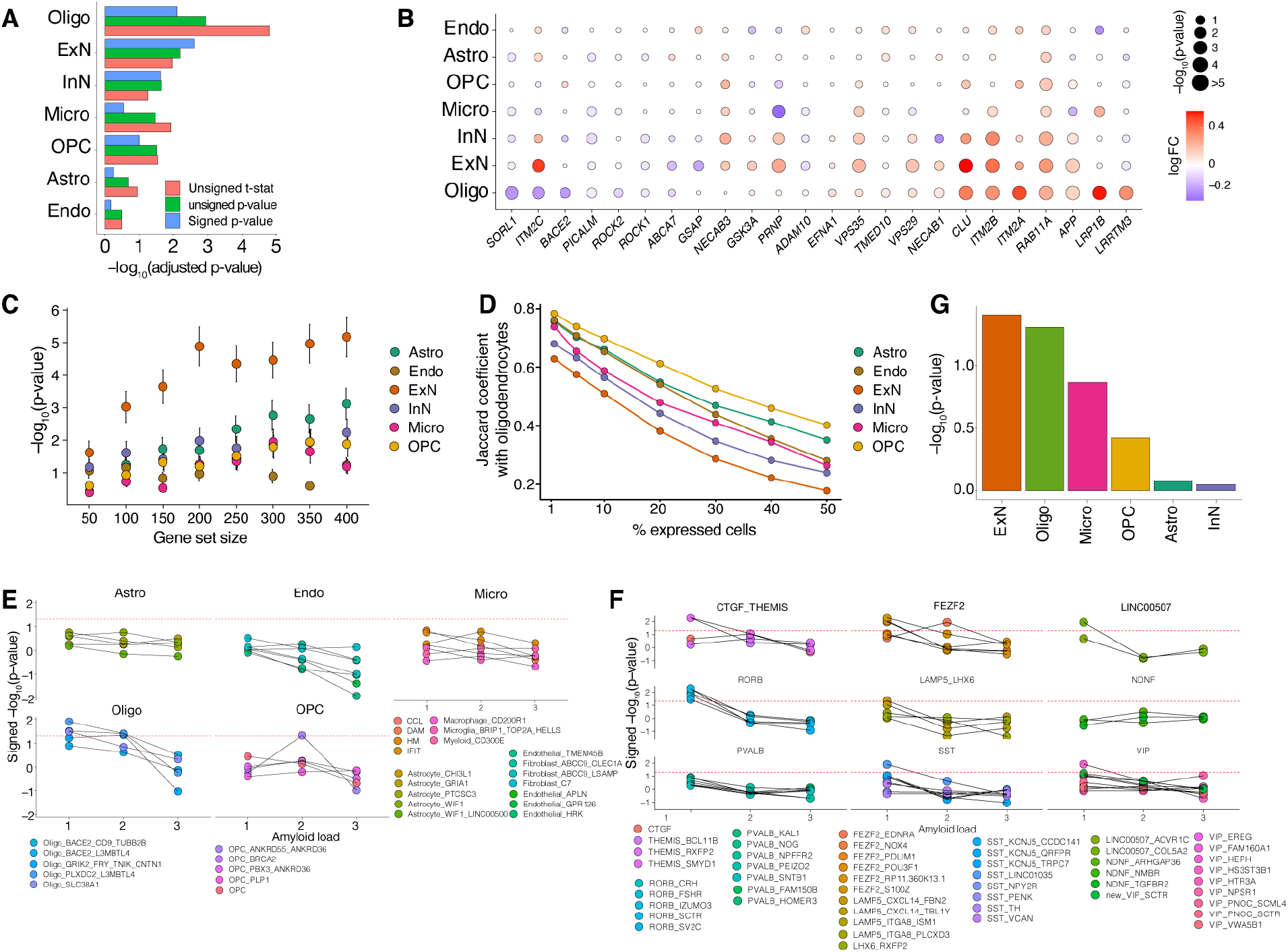
Nomination of amyloid-producing cell types in the human frontal cortex. ***A)*** *Bar chart showing -log_10_-transformed p-values for various ordering statistics (indicated at bottom right) based on GSEA of Aꞵ production and secretion geneset (**Table S7**) and cell class DE genes. **B)** Dot plot denoting differential expression of leading edge genes, identified by the GSEA of Aꞵ gene set in* Figure 5A*, in Aꞵ+ subjects versus Aꞵ-free biopsy samples. Color correlates with log-fold change and size correlates to -log_10_-transformed p-values. **C)** Correlation analysis results obtained via fGSEA (see **Methods**) comparing differentially expressed gene lists between all major cell classes and oligodendrocytes, with increasingly liberal thresholds (larger gene lists) for assigning univariate significance. **D)** Overlap of genes expressed in oligodendrocytes with other major cell classes. Different thresholds were selected to consider a gene as expressed (x-axis) based on the percentage of the cells in which the gene has non-zero UMI. **E,F)** Signed -log_10_-transformed p-values associated with fGSEA enrichment across increasing Aꞵ and tau burdens for all major cell types (glia in **(E)** and neurons in **(F)**) in human frontal cortex from DE analysis of the human biopsy dataset. **G)** Meta-analysis p-values for fGSEA of the Aꞵ associated gene set in the DE genes of two postmortem AD case-control datasets* ^3, 4^*, for six cell classes*.

**Figure S10.**
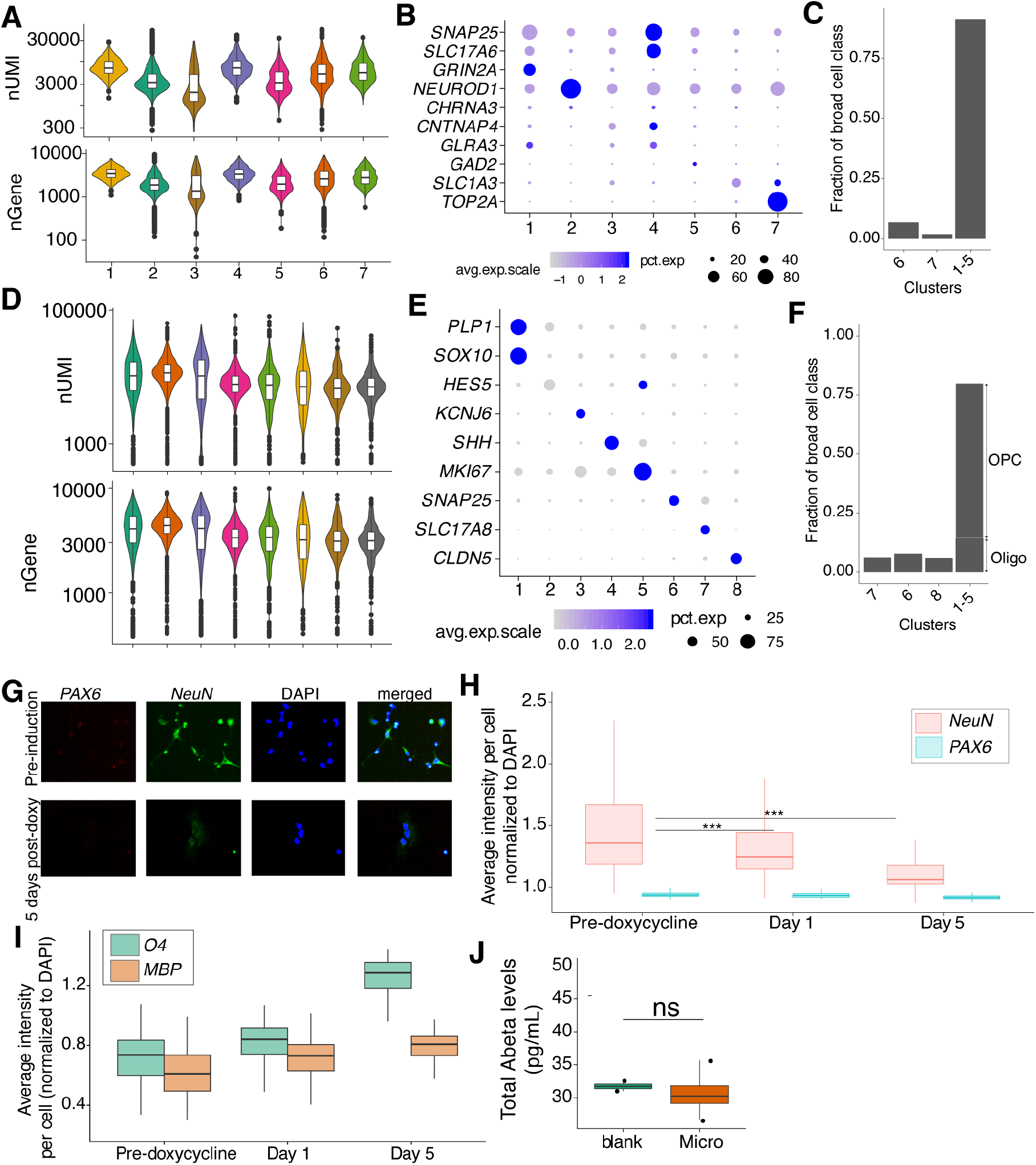
Single-cell transcriptomics and immunohistochemistry of ESC-derived oligodendrocyte and neuron cultures. ***A,D)*** *Violin plot of number of nUMI (top) and nGene (bottom) per cell type identified in single-cell transcriptomics of ESC-derived iExN (**A**) and (**D**) iOligo lineage cultures. In boxplots, center line, median; box limits, upper and lower quartiles; whiskers, 1.5x interquartile range; points, outliers. **B,E)** Key marker genes for cell types identified from single-cell transcriptomics of ESC-derived iExN (**B**) and (**E**) iOligo cultures. **C,F)** Composition of ESC-derived iExN (**C**) and iOligo (**F**) cultures based upon single-cell cluster annotations. **G)** Representative images of immunofluorescence stains of PAX6 and NeuN (**G**) in ESC-derived iOligo cultures. **H,I)** Box plots of average intensity values per cell (normalized to DAPI intensity) across days of differentiation for NeuN, PAX6 (**H**), O4, and MBP (**I**). Box upper and lower bounds represent upper and lower quartiles and Whisker distance from upper and lower hinges represents ≤1.5 times the interquartile range. Center line indicates the median value. **J)** Total Aꞵ levels from conditioned media isolated from ESC-derived microglia and blank control derived from unconditioned oligodendrocyte differentiation media. *** = p < 0.001, ** = p < 0.01, * = p < 0.05, NS = not significant. Statistical tests are based on a linear mixed-effect model for comparing immunofluorescence signal intensity per cell using each sample well as the levels of the random effect. Statistical significance for comparing amyloid beta protein values was determined via the Student’s t-test*.

## Notes

### Competing Interest Statement

The authors have declared no competing interest.

